# De novo design of small-molecule–induced conformational change

**DOI:** 10.64898/2026.08.03.742366

**Authors:** Jeffrey Chang, Nicholas F. Polizzi

## Abstract

Many biological proteins function by changing shape upon small-molecule binding. Here, we present a general strategy for designing de novo proteins that undergo small-molecule-induced conformational change. Our approach converts a preorganized small-molecule binding protein into a ligand-responsive shape-changer by adding a mobile lid domain that closes behind the ligand upon binding. Using this strategy, we converted an exatecan-binding protein into a drug-induced conformational switch. The lidded proteins showed considerably stronger binding affinity in the sub-nanomolar regime, 100-fold greater specificity to exatecan over a similar molecule, and ligand residence times up to several months, with tunable binding kinetics. We turned one design into a genetically encodable fluorescent biosensor of the drug, enabling potential clinical applications. Our results open the door to programming complex molecular function using vast chemical space.

## Main text

Biological proteins bind and respond to small molecules as essential aspects of their function. Current de novo design methods can reliably produce proteins that bind diverse small molecules^1– 8^, but eliciting a subsequent response remains an unsolved challenge^9,10^. The design of ligand-responsive proteins would extend the realm of complex molecular function beyond the scope of natural proteins. However, this design task is challenging because it contradicts the very principle that underlies the success of binder-design algorithms: preorganization^2,5,8^. Preorganized binders are designed to adopt identical structures with and without the bound ligand to minimize the loss of conformational entropy upon binding. While this paradigm can now readily produce binders with fast on-rates^7,8,11^, the off-rates are often also fast^5,7,11^, limiting achievable affinity. Furthermore, preorganization is incompatible with ligand-induced conformational change. Still, we wondered if we could build upon a static binder to elicit small-molecule response, while retaining the advantages of preorganization. Here, we implement a strategy used by natural proteins, such as streptavidin^12,13^ and Abl kinase^14,15^ (Fig. 1A), to add a mobile lid that confers small-molecule-induced shape change to an otherwise-preorganized binder (Fig. 1B). This general lid-based strategy produces de novo small-molecule-binding proteins endowed with sub-nanomolar affinities, exquisite selectivity, tunable on- and off-rates, and compatibility with spectroscopic readouts like FRET, unlocking a plethora of downstream applications in drug delivery, cell signaling, and biosensing.

**Fig. 1.**
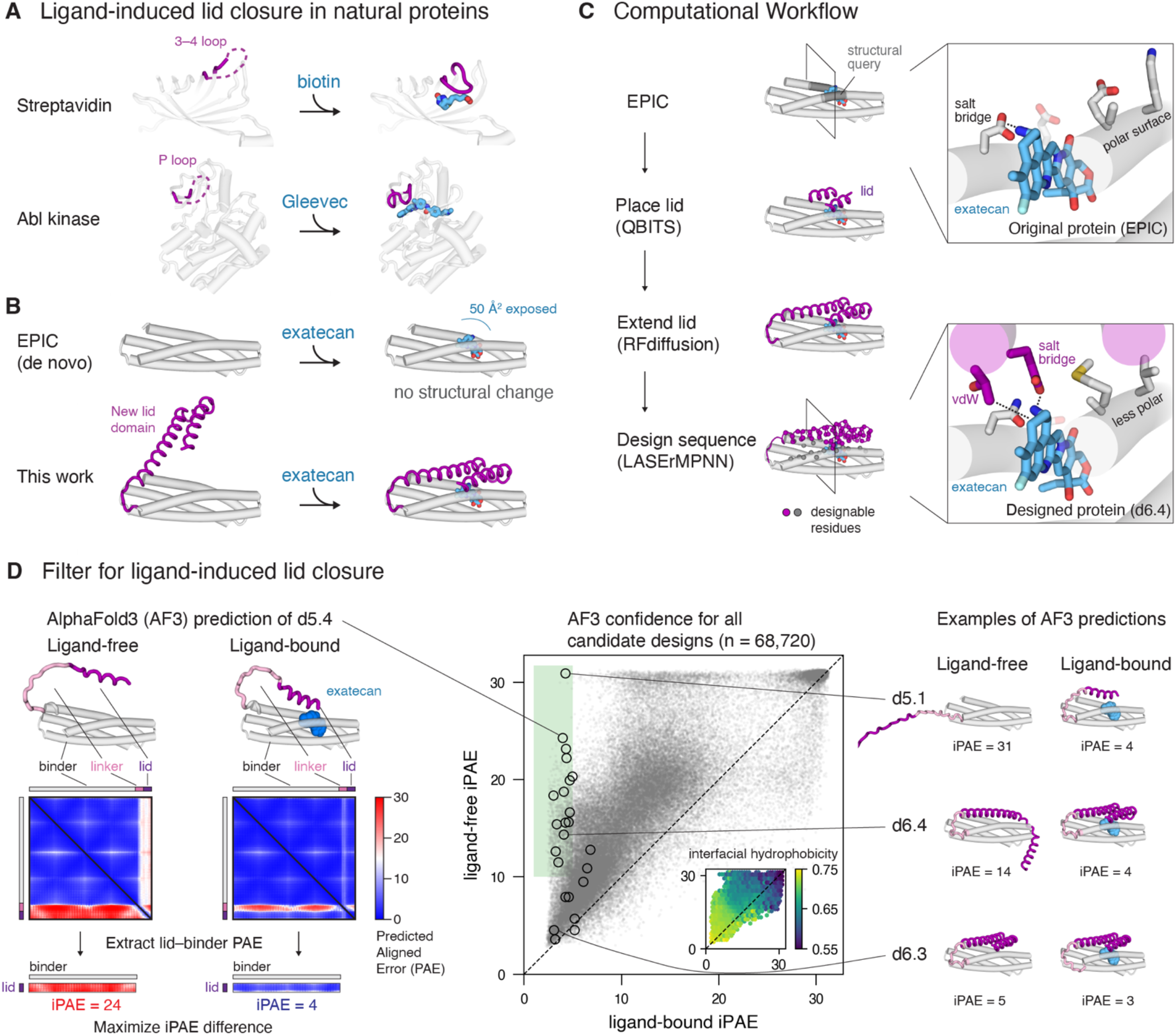
Strategy for de novo design of small-molecule-induced lid closure. (**A**) Examples of high-affinity natural proteins that use a lid (purple) that closes behind a ligand (blue) after binding. Apo/holo PDB codes: streptavidin-1swb, 1stp; Abl kinase-2hz4, 2hyy. (**B**) Schematic of the design goal to convert the de novo protein, EPIC, into a lidded binder. (**C**) Computational workflow for design of a lid domain fused to EPIC. Lid placement is seeded using QBITS and a two-helical search query (dark gray), extended with RFdiffusion, and joined to the binder domain with a flexible linker. The residues of the lid and the lid–binder interface (spheres) are designed using LASErMPNN. Insets: Example of a designed lid–binder interface, comparing the original protein EPIC (top) to the newly designed interface (bottom), showing the surface-exposed binder residues become less polar and the lid forms novel polar and apolar interactions with exatecan (blue). (**D**) Designed sequences are filtered using AlphaFold3 (AF3) predictions of ligand-bound and ligand-free proteins. As a proxy for ligand-induced lid closure, we select designs where, without ligand, AF3 is uncertain about the lid placement relative to the binder domain (high interfacial predicted aligned error, iPAE; left), but, with ligand, AF3 is confident in the lid placement (low iPAE). iPAE is calculated by averaging between all pairs of residues across the lid–binder interface. Center: scatterplot of iPAE for candidate designs (n = 68,720). The 24 designs selected for experimental validation are circled. Points in the green region represent favorable candidate designs with ligand-bound PAE < 5 (see Supplementary Note S1 for additional filtering criteria). Inset: Same 68,720 points of the larger iPAE scatterplot re-colored by hydrophobicity of the lid–binder interface, defined as the fraction of total interfacial surface area formed by non-polar atoms (higher number and yellower color is more hydrophobic interface).

We started from the de novo binder, EPIC, which buries the small-molecule drug, exatecan, within a highly preorganized four-helix bundle^8^. Most of the drug is encapsulated within rigid helices, but a small portion (50 Å^2^) protrudes from the binding site, as is common in de novo designed binders (Fig. 1B). We reasoned that exatecan’s exposed region, which includes a charged amine, could be leveraged to interact with an additional mobile “lid” domain that changes shape upon drug binding. Through newly designed drug–lid interactions, binding of the drug to the static “binder” domain of EPIC would drive lid closure (Fig. 1B). A protein designed to undergo exatecan-induced conformational change could be further engineered into a clinically relevant biosensor, since exatecan is a popular payload on antibody–drug conjugates. There is no known natural protein that can serve as a starting point for biosensor engineering, since the drug’s target is a protein–DNA complex with Topoisomerase I. EPIC provides an excellent starting point because it is small (17 K_d_a), stable (Tm > 95 °C), and its dissociation constant (K_d_ = 1.2 nM) is already in the clinically observed concentration range^16,17^.

While many natural proteins use flexible loops as lid domains, most de novo small-molecule binders house a ligand within rigid elements of secondary structure with little loop involvement^2,3,5,7,8,18^. We reasoned we could functionally recreate the features of a dynamic loop via a well-placed helix or pair of helices connected to the original binder by a flexible linker. For binding of exatecan to induce lid closure, the helical lid domain must interact favorably with the exposed portion of exatecan. Forming interactions with such a small target also requires the lid to interact with nearby surface residues—but not so strongly as to induce lid closure in the absence of drug. The lid of the drug-free protein must open to allow for ligand entry, which necessitates negative design.

There are a number of ways to achieve an open lid via negative design without explicitly defining its structure. For example, we reasoned that placing a negatively charged residue at the binder–lid interface could be effective. In the presence of drug, this residue would form a buried salt bridge with the drug’s positively charged amine; in the drug’s absence, lid closure would result in burial of an uncompensated negative charge, incurring a substantial desolvation penalty. Such positive- and negative-design strategies could be specified explicitly in the design algorithm, or they could be achieved implicitly by sampling many diverse sequences and selecting those predicted to satisfy a closed structure with drug and an open structure sans drug.

To place a new lid domain with a good packing geometry near the protrusion of exatecan in EPIC, we searched the protein data bank (PDB) for proteins containing a similar structural motif as the two helical segments flanking the ligand [Fig. 1C, dark gray, Cα root mean square deviation (RMSD) < 1.2 Å]. We wrote a structural bioinformatics algorithm called QBITS (query-based interfacial tertiary structures) to identify common interfacial geometries among the matching proteins. QBITS nominated the most prevalent interfaces as initial placements for a helical lid domain (supplementary text S1). These initial placements were extended using RFDiffusion^19^ and joined to the binding domain with a flexible (GGS)n linker (n = 4–7). Our sequence-agnostic approach gave a few hundred single- and double-helix placements for subsequent sequence design with the graph neural network, LASErMPNN^8^.

Using LASErMPNN, we designed the sequence of the lid and the residues of EPIC’s 4-helix bundle at the lid interface (Fig. 1C, spheres). While we filtered for sequences that formed a salt bridge between the lid and exatecan’s amine, we wanted to employ a ranking procedure that was agnostic to any specific, user-defined intent. We therefore ranked the resulting ∼60,000 sequences of candidate 5-helix and 6-helix designs based on AlphaFold3^20^ (AF3) predictions of exatecan-free and exatecan-bound structures (Fig. 1D). We aimed to find designs with low interfacial predicted aligned error (iPAE) between the lid and binder domains in the bound state, and high iPAE between them in the unbound state. In such designs, AF3 is confident in the lid placement only when the ligand is present.

We assessed the pairs of predictions by their difference in iPAE and lid RMSD and found the largest differences occurred with interfacial sequences of moderate polarity, agreeing with our biophysical intuition. Lids that were dubiously predicted for both the ligand-free and ligand-bound forms were too polar, and lids that were confidently closed were too hydrophobic (Fig. 1D inset). For lids of moderate polarity, the flexible (GGS)n linker between the binder and lid domains challenges AF3 to predict the correct helical packing register of the lid–binder interface. In the best-ranked designs, we found that the lid–ligand salt bridge, which is formed only when the ligand is present, locks in the intended helical register of the lid; in the absence of ligand, the lid is predicted to be either fully dissociated or register-shifted, with a larger iPAE (Fig. 1D). We selected 24 designs with large ligand-dependent change in iPAE and lid RMSD, including some designs with low iPAE in both ligand-bound and ligand-free forms to test the effectiveness of our negative design strategy (tables S1–S3).

### Designs show sub-nanomolar affinity and increased specificity

We ordered synthetic DNA for the 24 designs and expressed them in *E. coli*. All the lidded proteins expressed in good yield and still bound exatecan with K_d_ < 1 µM (table S4). To assess the designs for exatecan-induced lid closure, we covalently attached a fluorescein dye at a solvent-exposed position of the lid (via a Cys mutation) and measured its fluorescence anisotropy (Fig. 2A). Upon addition of excess exatecan, 4 designs exhibited a marked and reproducible increase in fluorescence anisotropy (Fig. 2A), indicating some degree of lid closure (supplementary text S2). As a more sensitive measurement of exatecan-dependent lid closure, we compared the rate of tryptic cleavage of the lid in the presence and absence of exatecan (Fig. 2B, fig. S1); this limited proteolysis assay identified 6 additional designs as weaker hits earmarked for further negative design. Most design hits had ΔiPAE > 3 (3/4 from anisotropy and 6/9 from limited proteolysis, fig. S2), with some hits, such as d6.4, nearing ΔiPAE of 10. We focused on the top design nominated by each assay—d6.9 by fluorescence anisotropy and d6.4 by partial proteolysis—for further characterization. Both d6.4 and d6.9 were identified as hits in both assays, monomeric by size exclusion chromatography (with and without ligand, fig. S3), and predicted to completely enclose exatecan via lid-bestowed hydrophobic and polar interactions, with no exposed ligand surface area (0 Å^2^).

**Fig. 2.**
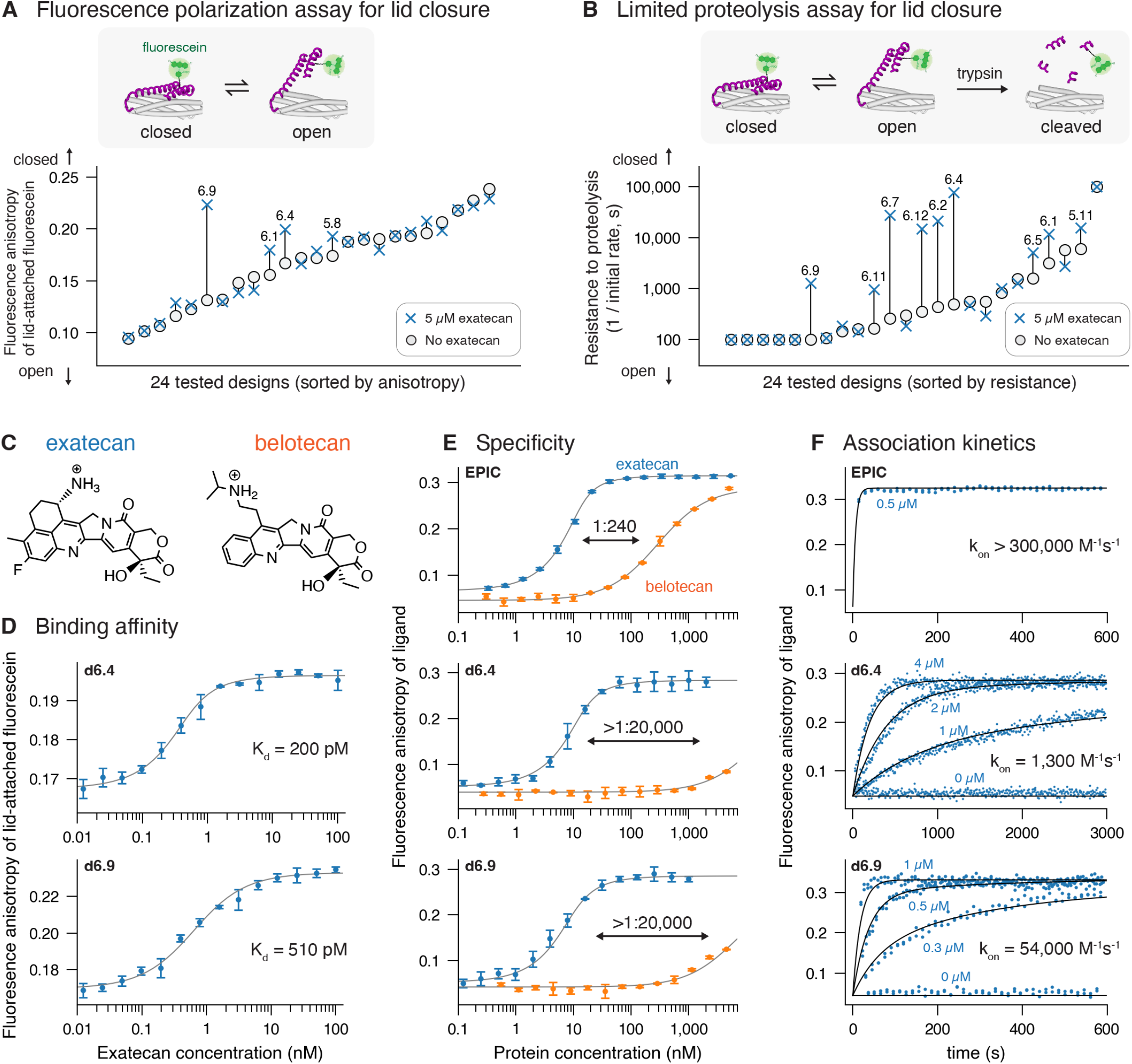
Designs show ligand-induced lid closure and improved binding. (**A**) Above: schematic of fluorescence polarization assay for exatecan-induced lid closure. Fluorescein is covalently attached to the lid through a solvent-exposed cysteine, and its fluorescence anisotropy is measured at equilibrium to assess lid open *vs* closed populations. Below: the 24 designs were tested for exatecan-induced lid closure by measuring fluorescence anisotropy of fluorescein-labeled protein (75 nM) without (gray circles) and with exatecan (5 µM, blue crosses) in PBS pH 6.5 (0.01% w/v PEG 3350). For each design, the exatecan-induced difference in anisotropy is represented with a vertical line. Designs with change in anisotropy > 0.015 are labeled by name. (**B**) The 24 designs were also tested for exatecan-induced lid closure by comparing proteolytic resistance in the presence (blue crosses, 5 µM) and absence (gray circles) of exatecan. Fluorescein-labeled proteins were treated with trypsin (0.1 mg/mL), and the initial rate of lid cleavage was measured via the time-dependent decrease in anisotropy of the lid-attached fluorescein. A greater resistance to proteolysis (slower rate) indicates a smaller subpopulation in a protease-accessible open conformation. Designs where adding exatecan caused a >2-fold change in proteolytic resistance are labeled by name. (**C**) Chemical structures of exatecan and belotecan. (**D**) Binding titration of exatecan to d6.4 and d6.9, monitoring the change in anisotropy of lid-attached fluorescein ([protein] = 333 pM). K_d_ values are derived from global fits across three protein concentrations (fig. S3). (**E**) Binding titration of EPIC, d6.4, and d6.9 with constant amount of exatecan (blue) and belotecan (orange), monitoring the intrinsic fluorescence of the ligand ([ligand] = 10 nM). A fit to a quadratic binding curve is shown (K_d_ = 1.2 nM, < 2 nM, < 2 nM with exatecan; K_d_ = 290 nM, > 10 µM, and > 10 µM with belotecan; values for EPIC, d6.4, and d6.9, respectively). Note that K_d_ < 2 nM cannot be distinguished in this experiment due to [ligand] = 10 nM. (**F**) Association kinetics of exatecan to EPIC, d6.4, and d6.9, at various protein concentrations (blue labels), based on the intrinsic fluorescence anisotropy of the ligand ([exatecan] = 50 nM). The rate constant was fit globally across several protein concentrations (fig. S8). Error bars show standard deviation across 4 (**D**) or 3 (**E**) technical replicates.

Binding of d6.4 and d6.9 to exatecan was too tight to measure accurately by monitoring exatecan’s weak intrinsic fluorescence. Instead, we measured the fluorescence anisotropy of the bright fluorescein label attached to the lid as a function of titrated exatecan (Fig. 2D). Using this more sensitive approach to reduce protein concentration, we measured dissociation constants for d6.4 and d6.9 in the sub-nanomolar range (200 ± 30 pM and 510 ± 50 pM from global fitting, fig. S4; error bars represent standard errors of fits). The enhanced binding affinity of the lidded proteins relative to EPIC (K_d_ = 1.2 nM) suggested that the lid formed favorable interactions with exatecan to slow its dissociation. Indeed, we inferred lid–ligand interaction energies of ΔΔG = –3.2 and – 2.3 kcal/mol for d6.4 and d6.9, respectively, by using partial proteolysis rates to quantify the shift in the lid’s conformational equilibrium (260- and 50-fold, respectively, fig. S5, see supplementary text S3).

To determine whether lid–ligand interactions also conferred a greater specificity for exatecan over similar compounds, we measured the binding affinity of d6.4 and d6.9 towards belotecan, a tecan analog which differs from exatecan most prominently near the amine functional group (Fig. 2C). While the parent protein EPIC binds belotecan with K_d_ = 290 nM (1:240 specificity ratio), we found that the lidded proteins bound belotecan far more weakly, with K_d_ > 10 µM, corresponding to a specificity ratio of >1:20,000 (Fig. 2E). The lidded designs were thus >100-fold more specific than the lidless original with respect to discerning between highly similar ligands.

To gain more insight into the high affinity and specificity of d6.4 and d6.9, we solved crystal structures of both proteins bound to exatecan (Fig. 3, fig. S6, tables S5 and S6). Both structures agreed closely with their AF3 predictions (Cα RMSD 0.3 Å and 0.4 Å, ligand heavy-atom RMSD 0.6 Å and 1.2 Å, respectively) and showed that the lid is correctly placed in the designed position and helical register (Cα RMSD 1.1 Å and 1.0 Å, respectively). The two proteins were designed with the same main-chain placement of the lid but with different side-chain interactions at the binder–lid interface (68% sequence identity of designable residues, fig. S7). In both structures, a designed salt bridge between exatecan’s amine and the Glu41 sidechain is clearly resolved (Fig. 3B,C), along with hydrophobic packing of the Thr44 sidechain with a portion of exatecan that is solvent-exposed in EPIC. Notably, d6.9 buries an additional Glu residue at position 93, whereas this residue is a Met in d6.4. As a result, d6.9 shows more structured waters at the binder–lid interface (Fig. 3C). Overall, in both d6.4 and d6.9, exatecan is completely buried with no exposed surface area (0 Å^2^) because the lid covers the binding site. In contrast to EPIC, the lidded proteins leverage the entire surface of the ligand for molecular recognition.

**Fig. 3.**
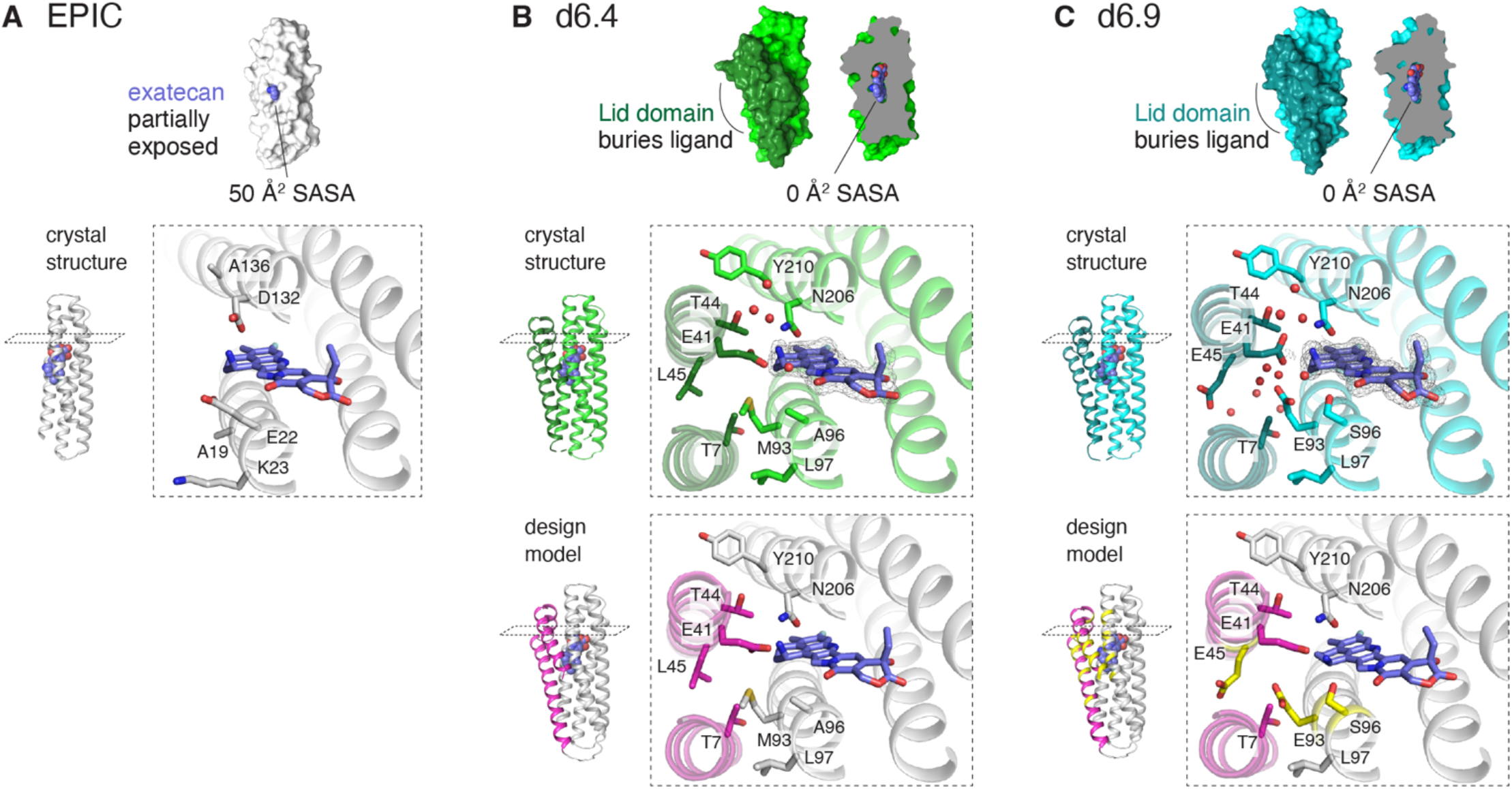
Crystal structures of EPIC and lidded designs d6.4 and d6.9. (**A**) Crystal structure of exatecan-bound EPIC, the starting point for lid design. Above, the structure of EPIC shown in surface representation, with solvent-accessible surface area (SASA) of exatecan (blue) indicated. Inset, sidechains of some residues near the solvent-exposed region of exatecan that were redesigned to accommodate the lid. (**B**) Crystal structure of d6.4 bound to exatecan. The cutaway illustrates that exatecan is completely buried by the new lid domain, with no solvent-exposed ligand surface. The insets show residues at the newly-designed lid–binder interface with side-chain conformations in near-complete agreement with the AlphaFold3 prediction (below, Cα RMSD = 0.3Å). The electron density of exatecan is shown with a grey mesh (composite omit 2mF_o_ − DF_c_ map, 1σ), and crystallographically resolved water molecules are shown as red spheres. (**C**) Crystal structure of d6.9 bound to exatecan (above) compared to the AlphaFold3 prediction (below, Cα RMSD = 0.4Å). Residue identities that differ between d6.4 and d6.9 (29 residues total) are colored in yellow (fig. S7). Design d6.9 shows a larger number of resolved waters in the interface (red spheres).

Since the lid completely occludes the drug’s binding site in d6.4 and d6.9, the lid must open, at least partially, to allow for the drug’s entry or egress. To measure the kinetics of drug entry, we monitored the time-dependent increase in fluorescence anisotropy of exatecan after mixing with protein. Designs d6.4 and d6.9 bound exatecan with on-rates of 1,300 M^-1^s^-1^ and 54,000 M^-1^s^-1^, respectively (Fig. 2F, fig. S8). Similar rates were observed in a separate experiment that tracked the anisotropy of lid-attached dye (fig. S9). Although slower than EPIC, these on-rates spanned a wide range that suggested they could be tuned. Using the equation k_off_ = K_d_ × k_on_, we inferred off-rates of 2×10^-7^ s^-1^ and 3×10^-5^ s^-1^ for d6.4 and d6.9, respectively. The off-rate of d6.4 is extremely slow (1/k_off_ ∼ 2 months), making the protein–drug complex pseudo-covalent. While slow off-rates are desirable for therapeutic applications that involve slow release of drug from a circulating drug– protein complex, sensing applications often demand faster kinetics^21^. We therefore set out to tune the binding kinetics of d6.4 to tailor it toward biosensing.

### Rational tuning of conformational equilibrium and binding kinetics

The slow on-rate of d6.4, together with the high drug-free equilibrium anisotropy of the dye label, strongly suggested the lid was partially associating with the 4-helix bundle in the absence of ligand. We therefore sought an approach to further open drug-free d6.4 while maintaining closure of drug-bound d6.4. We hypothesized that destabilizing the lid–binder interface would shift the conformational equilibrium of drug-free d6.4 towards a more open form, thereby accelerating the on-rate and increasing the magnitude of the conformational shift upon binding, which we could ultimately link to a genetically encodable FRET readout. We considered two strategies for destabilizing the interface: hydrophobic-to-polar substitutions or lid truncation.

Comparison of d6.4 to the more open d6.9 immediately suggested that a Met^93^→Glu^93^ mutation could sufficiently destabilize the binder–lid interface. This substitution introduces polar sidechain atoms that would require energetically costly desolvation (and potentially protonation) upon lid closure. The same M93E substitution could also be identified by comparing LASErMPNN-predicted amino-acid probabilities of open and closed d6.4 (fig. S10). LASErMPNN predicts Glu93 has a small but considerable probability in the closed state and substantially higher probability in the open state. We cloned and purified d6.4^M93E^ from *E. coli*. As predicted, drug-free d6.4^M93E^ showed a decreased equilibrium fluorescence anisotropy of lid-attached dye, which more closely matched that of an always-open control protein (Fig. 4B). Importantly, upon binding exatecan, d6.4^M93E^ showed the same high anisotropy as unsubstituted d6.4. In step with the equilibrium results, the measured on-rate increased 30-fold to 37,000 M^-1^s^-1^, without appreciably affecting the binding affinity (K_d_ = 100 ± 10 pM, Fig. 4C, fig. S4). Thus, a single polar mutation in the interface can sufficiently shift the lid’s conformational equilibrium and quicken the on-rate.

**Fig. 4.**
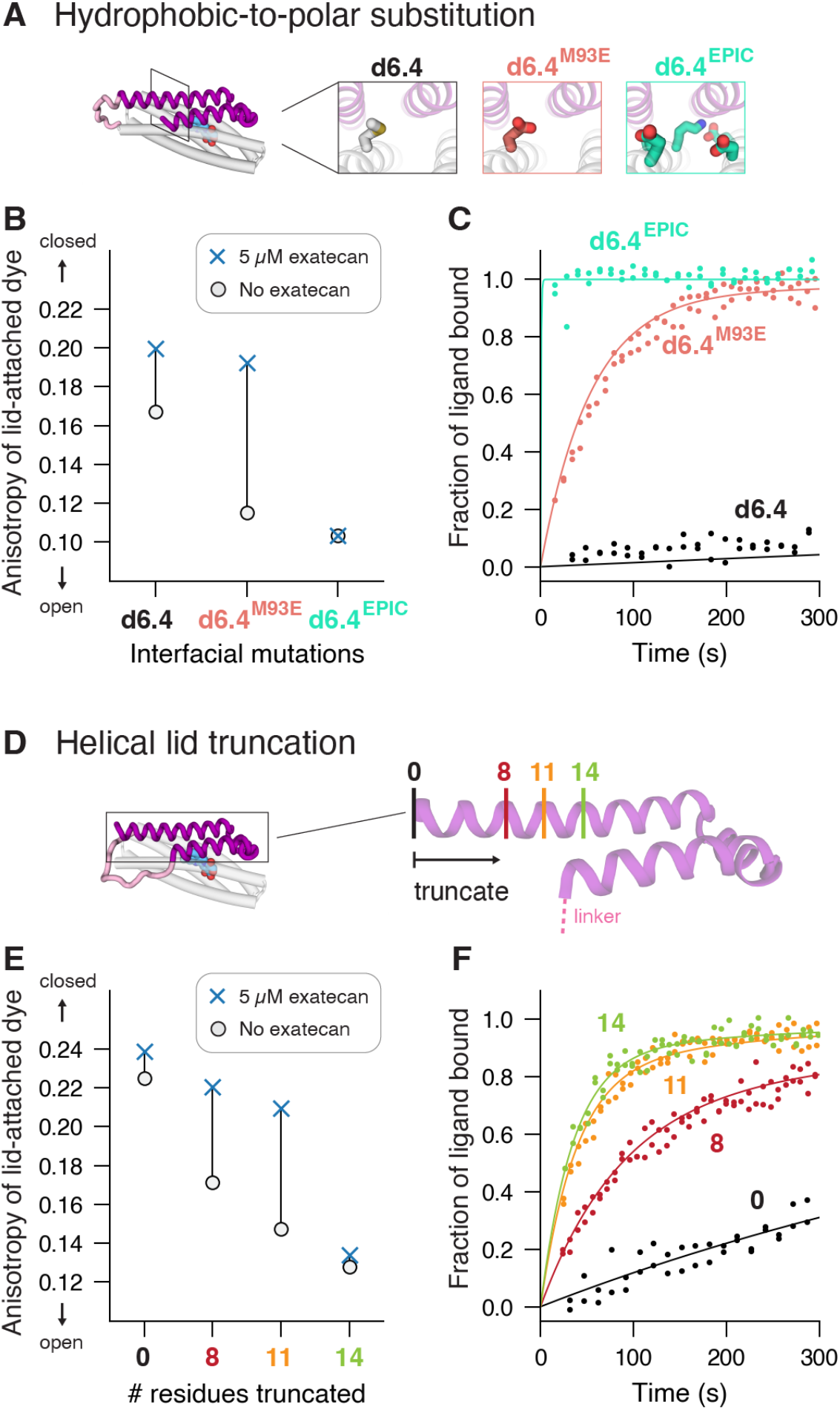
Rational tuning of d6.4 conformational equilibrium and binding kinetics. (**A**) Mutation series of d6.4: the original d6.4 (black); the single hydrophobic-to-polar substitution Met^93^→Glu^93^ (d6.4^M93E^, salmon color); and a full reversion of the binder domain to the sequence from EPIC (d6.4^EPIC^, mint color), which is expected to disrupt lid closure, leading to an open lid conformation. (**B**) Ligand-induced lid closure of the mutant series as measured by the anisotropy of lid-attached fluorescein in d6.4, d6.4^M56E^, and d6.4^EPIC^ (75 nM labeled protein). The vertical line represents the difference in anisotropy between exatecan-bound (blue x, saturating [exatecan] = 5 µM) and exatecan-free (gray o) conditions. (**C**) Association kinetics of exatecan to d6.4, d6.4^M56E^, and d6.4^EPIC^ ([unlabeled protein] = 500 nM, [exatecan] = 25 nM, PBS pH 6.5). The fraction bound is determined from fitting the intrinsic fluorescence anisotropy of exatecan. Curves are derived from a global fit to a kinetic model described in fig. S8. (**D**) A series of truncations, which remove 0, 8, 11, or 14 residues from the C-terminus of circularly-permuted d6.4, systematically tunes interface strength. (**E**) and (**F**), same conditions for the truncation series as (**B**) and (**C**). These results are summarized in table S7.

Because buried surface area at a protein–protein interface correlates with affinity between proteins^22^, we reasoned that another systematic way to tune the lid conformational equilibrium— and therefore the on-rate—would be to truncate the lid’s helix by varying amounts. The terminus of d6.4’s lid domain is positioned close to the ligand; so, to avoid perturbing lid–ligand interactions, we moved the terminus via circular permutation (Fig. 4D, fig. S11). This permutation also brought the N- and C-termini closer together to potentially maximize a FRET signal (see below). A series of lid truncations showed that an 11-residue deletion from the C-terminus gave the optimal exatecan-dependent change in anisotropy of the lid-attached dye (Fig. 4E). The smaller interface of the deletion mutant, d6.4^cpΔ11^, opened the drug-free protein (lowered the dye anisotropy) to a similar degree as the M93E mutation, while still maintaining good closure upon drug binding (similarly high dye anisotropy). Unlike the polar substitution, we found that truncation of the lid caused the binding affinity to become slightly weaker (K_d_ = 4.4 ± 0.7 nM, fig. S4). As expected, d6.4^cpΔ11^ showed more rapid binding kinetics than d6.4, and overall, the on-rate increased monotonically with the size of the deletion (from 10,000 to 57,000 M^-1^s^-1^, Fig. 4F, fig. S8). Truncation of the helical lid thus presents a simple and general way to tune the lid’s conformational equilibrium and modulate binding kinetics.

### Ligand-induced lid closure enables FRET sensing

With faster binding kinetics in hand, we sought to transform the circularly permuted d6.4^cpΔ11^, which has its N- and C-termini in close proximity (fig. S11), into a genetically encodable FRET sensor of exatecan. A change in FRET upon drug binding not only corroborates a ligand-induced conformational change but also demonstrates that lid closure can be coupled to a downstream function that enables cellular experiments or clinical monitoring of the drug. De novo design of biosensors has seen tremendous recent progress but remains challenging, often requiring screening of thousands or more designs^5,7,23–26^. We wondered if the relatively simplistic architecture of d6.4^cpΔ11^ would be more amenable to rational engineering.

Because the mobile lid is positioned at one terminus and the static binder at the other, we reasoned that fusing the fluorescent proteins, mTurquoise2 (mT) and mVenus (mV), to the two termini of d6.4^cpΔ11^ would couple exatecan-induced lid closure to a change in FRET (Fig. 5A). We constructed the fusion protein, mT-d6.4^cpΔ11^-mV, and found the donor-to-acceptor peak ratio increased substantially upon addition of exatecan (0.79 to 1.13, Fig. 5B). By measuring change in FRET while titrating exatecan, we determined a low-nanomolar binding affinity (K_d_ = 1.5 ± 0.3 nM from a global fit, fig. S12), which should provide a suitable dynamic range for detecting clinically relevant concentrations of the drug. To determine if competing metabolites would interfere with FRET-based detection of exatecan, we tested mT-d6.4^cpΔ11^-mV with varying amounts of exatecan spiked into human saliva, urine, and serum. We observed a similar FRET change to that in phosphate-buffered saline (Fig. 5C and 5D), implying that lid closure is specific to exatecan, even within complex biological environments. These results show promise for the potential use of de novo biosensors in clinical settings.

**Fig. 5.**
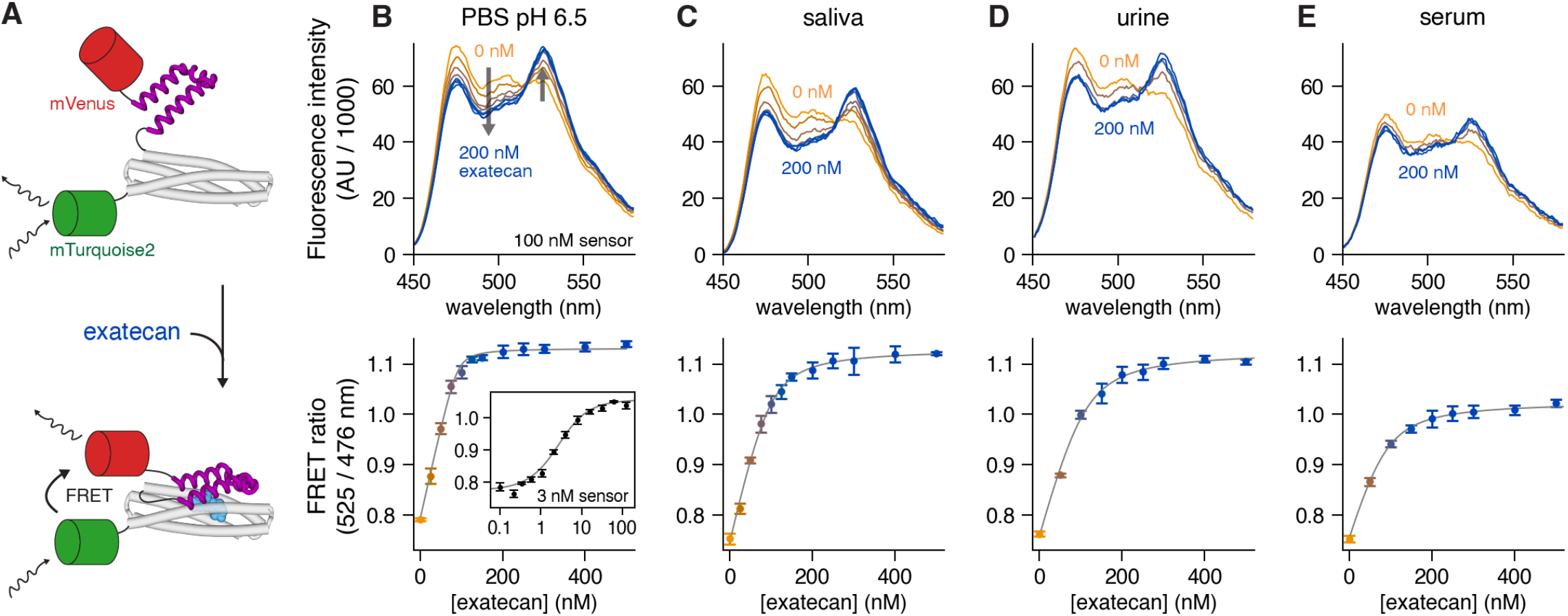
Ligand-induced lid closure enables FRET sensing in complex biological environments. (**A**) Schematic of a de novo FRET sensor of exatecan via fusion of fluorescent proteins, mTurquoise2 and mVenus, to the termini of d6.4^cpΔ11^. Exatecan-induced lid closure increases FRET efficiency. (**B**) Above, emission spectra of the FRET sensor at varying exatecan concentrations in PBS pH 6.5, with a fixed sensor concentration of 100 nM. Below, the donor-to-acceptor peak ratio (525 nm / 476 nm) is plotted as a function of exatecan concentration. Error bars show standard deviation of 4 technical replicates. The gray curve is a fit to a 1:1 binding model with K_d_ = 1.5 nM (with K_d_ derived from a separate global fit, fig. S12). Inset, FRET ratio as a function of exatecan concentration, with the sensor concentration fixed at 3 nM. (**C, D, E**) The exatecan sensor is functional in saliva, urine, and serum. Exatecan was spiked at 2x the indicated concentration into the biological environment and mixed 1:1 with 200 nM of sensor in PBS pH 6.5 to a final sensor concentration of 100 nM. Background fluorescence from the environment is subtracted before plotting spectra and calculating peak ratios. The fit gray curve includes an additional fitting parameter to account for the binding of exatecan to environmental proteins such as albumin in serum (K_d_ = 43 µM to exatecan^8^), which causes the saturation point of the titration curve to be shifted rightward from the 1:1 point at 100 nM. The low FRET efficiency at [exatecan] = 0 nM across PBS and each of the biological matrices indicates that environmental metabolites do not induce lid closure of the sensor.

## Discussion

The design of proteins that bind and respond to small-molecule ligands has been a grand challenge in molecular engineering^9,10,24,27–29^. Here, we devised a general method to turn a static de novo small-molecule binder into a ligand-induced conformational switch. Our approach—inspired by biological examples of streptavidin^12,13^, GPCRs^30,31^, and Abl kinase^14,15^—adds a conformationally mobile lid that closes behind the ligand, consistent with an induced-fit pathway. In streptavidin, closure of the 3-4 loop drives high affinity with biotin^12,13,32^; in GPCRs like µ-opioid and serotonin receptors, closure of the ECL2 loop accounts for exceptionally slow drug dissociation^30,31^; in Abl kinase, closure of the P-loop enables the specificity with and long residence time of Gleevec^14,15^. Analogously, our de novo-designed lid closure achieved increased affinity and specificity to exatecan, as well as tunable on- and off-rates. The straightforward engineering of mT-d6.4^cpΔ11^-mV showed that lid closure can couple small-molecule binding to a functional readout such as FRET. The only design requirement is that part of the ligand is exposed to solvent (and therefore available for forming new ligand–lid interactions), which is readily achievable with current design algorithms^2,3,5,7,8^.

Intramolecular lid closure provides unique opportunities not afforded by other sensing strategies such as chemically induced dimerization (CID), which designs a second protein to conditionally bind the ligand-bound protein^23–26,33,34^. First, with intramolecular conformational change, the binding readout does not depend on titrating the appropriate concentration of a second protein, simplifying expression in living systems. Second, CID systems must overcome roto-translational entropy to form a ternary complex, which favors rigid preorganization of both binding partners and targeting large hydrophobic interfaces^23,25^. By contrast, tethering a mobile lid to a static binder pays the roto-translational entropy cost upfront and increases the lid’s effective concentration (> 1 mM)^35^. This allows targeting smaller, more polar epitopes of small molecules—which are often the most solvent-exposed portions in designed binders—using a ligand surface area much smaller than in known CIDs (50 Å^2^ in exatecan–EPIC *vs* 520 Å^2^ in rapamycin–FKBP, fig. S13). A tethered lid does not require strict preorganization and can explore myriad ligand-free conformations such as helical phase- or register-shifts, kinks, and disorder-to-order transitions. However, the requirement for negative design is stringent, since the affinity of the lid for the binder must be weak (K_d_ > 1 mM) in the absence of ligand to allow for ligand entry. (In this light, most designed CID pairs^5,25,36,37^, if tethered, would likely prematurely associate in the absence of the ligand, since ligand-free K_d_ is typically ∼10 µM.) We found that energetics of the lid–binder interface can be rationally tuned through hydrophobic-to-polar mutations or lid truncations. This tunability means designs can first be screened for even trace amounts of ligand-induced lid closure using a sensitive assay like limited proteolysis, and then readily optimized using the principles outlined here. Indeed, we achieved small-molecule-driven lid closure after testing only 24 designs, suggesting that rapid conversion of static binders to functional switches is now within reach of de novo design. This high success rate contrasts with recent CID design campaigns^5,23,25,34,37^, which required screening of thousands of designs followed by library-based optimization.

Our work offers a straightforward roadmap for designing ligand-induced conformational change. Lid closure provides a general mechanism for tuning binding kinetics, driving ultrahigh affinity, and enhancing specificity. Here, lid closure was driven by a tiny, polar surface presented by the bound ligand, a particularly challenging epitope to target. Thus, we expect our strategy to be applicable to a wide portion of chemical space, including clinically relevant drugs like exatecan. The ability to tackle polar surfaces and to choose predefined ligands across chemical space contrasts with previous work^38^, which relied on large, apolar surfaces and custom-made amphipathic helices as ligands. While natural switching motifs can be repurposed^10^, de novo lid design—built atop de novo small-molecule binders—could feasibly allow conformational change to be driven by potentially any small molecule, even those without obvious natural-protein starting points. Our work provides a recipe for designing novel ligand-induced shape change that can be readily coupled to a variety of downstream functions to enable the de novo design of complex biological systems.

## Methods

See supplementary information.

### Data Availability

Coordinates and data files for X-ray crystal structures of exatecan-free d5.11, exatecan-bound d5.11, exatecan-bound d6.4, and exatecan-bound d6.9 have been deposited in the PDB with accession codes 37RS, 37RT, 37RV, and 37RU, respectively. AlphaFold3 predictions for all 24 designs, exatecan-bound and exatecan-free, are deposited in Zenodo at https://doi.org/10.5281/zenodo.21780330.

### Code Availability

Open-source code for QBITS can be found in github at https://github.com/npolizzi/Qbits/releases/tag/v1.0.0.

## Acknowledgements

We thank Kaia Slaw and Franziska L. Sendker for assistance collecting diffraction datasets. We thank Harrison Wang and Steve G. Boxer for useful suggestions. We thank E. S. Fischer, Wesley P. Wong, Spencer C. Guo, Pranav V. Lalgudi, and Samuel P. Berry for helpful feedback on the manuscript. We thank SBGrid for support of on-site compute infrastructure and crystallography software. This research used resource 17-ID-1 of the National Synchrotron Light Source II, a U.S. Department of Energy (DOE) Office of Science User Facility operated for the DOE Office of Science by Brookhaven National Laboratory under Contract No. DE-SC0012704.

## Author contributions

N.P. conceptualized the idea. J.C. performed the computations and experiments. J.C. and N.F.P. analyzed data and wrote the paper.

## Materials and Methods

### DNA Cloning

All designed amino acid sequences were codon optimized for *E. coli* expression using the online codon optimization tool from GenScript. The following overhang sequences were added to the 5’ and 3’ ends as cloning sites for golden gate assembly with the BsaI cut site:

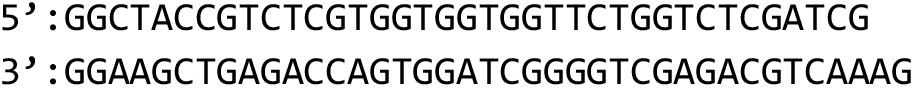

Synthetic genes were ordered as eblocks from Integrated DNA Technologies. Using the NEB Golden Gate assembly kit, the gene fragment was inserted into a custom expression plasmid, peAIP32 (a slight variant of Addgene #258140 with TCG inserted at position 472). The plasmid contains an N-terminal 10x His tag and a SUMO tag followed by a single extra serine residue as a cloning scar. The assembled product was transformed into chemically competent BL21-Gold (DE3) *E. coli* cells (Agilent 230132) using the *Mix & Go!* kit and protocol (Zymo T3001). The outgrowth was plated on LB-agar with 50 µg/mL kanamycin, and single colonies were picked for whole-plasmid nanopore sequencing. After verifying that the plasmid was correctly assembled with the correct insert sequence via long-read sequencing (Quintara Biosciences), a glycerol stock was prepared for subsequent protein purification.

### Protein Expression and Purification

Initial design screening and small-scale expression was carried out as 3 mL bacterial growths in 24-well plates. Growth was inoculated either with a fresh colony or with 3 µL (1:1000) of an overnight starter culture grown from the glycerol stock. The cells were grown in an autoinduction medium (Miller’s Luria Broth with 50 µg/mL kanamycin and 12.5 g/L lactose) containing lactose to induce protein expression in mid-log phase. The culture was incubated overnight (16–20 hr) at 30°C with 220 rpm shaking.

Following overnight growth, the cells were pelleted at 3,000 g for 10 minutes and then thoroughly resuspended in 300 µL of Lysis Buffer I containing: 10 mM phosphate buffer at pH 7.4, 2.7 mM KCl, 500 mM NaCl, 20 mM imidazole, 1 µg/mL DNAse I, a small dab of lyophilized lysozyme (Fisher AAJ6070103), and 1× BugBuster® Protein Extraction Reagent (Millipore 70921). After gentle shaking for 10 minutes, the cell debris was pelleted by centrifugation at 3,000 g for 30 minutes, and the supernatant was carefully transferred to a separate 24-well plate (Cytiva 7701-5102) containing 50 µL of a Ni-NTA magnetic bead slurry (GenScript L00295) equilibrated with Wash Buffer (10 mM phosphate buffer at pH 7.4, 2.7 mM KCl, 500 mM NaCl, 20 mM imidazole). The Ni-NTA magnetic beads were gently rocked for 30 minutes and then washed 5 times with 2 mL of Wash Buffer. In between wash steps, the beads were separated from the supernatant using an arrayed rack of 24 O-shaped neodymium magnets. To elute the proteins, the beads were first equilibrated into phosphate-buffered saline (PBS) pH 6.5 (Sigma P4417-100TAB) and then incubated with 1 µM His-tagged *Cth* Ulp1 protease (*1*) for 1–2 hours at 25 °C with mild shaking. The supernatant (containing the cleaved protein) was collected, leaving the SUMO tag and protease bound to the Ni-NTA beads. Purity was assessed by gel electrophoresis. The final purified protein was stored at 4 °C in PBS pH 6.5. Typical yields were 10–20 µM in 200 µL of eluted protein, corresponding to 20–40 mg protein per liter of expression culture.

For larger-scale protein preparations, expression cultures were grown under the same conditions in 500 mL of autoinduction media in a 4 L Erlenmeyer flask. The cells were resuspended in Lysis Buffer II (10 mM phosphate buffer at pH 7.4, 2.7 mM KCl, 500 mM NaCl, 20 mM imidazole, 1 µg/mL DNAse I) and were lysed by sonication on ice. The lysate was centrifuged at 11,600 g for 30 minutes to remove cell debris, and the supernatant was loaded onto a column of Ni-NTA agarose. The column was washed with 5 column volumes of Wash Buffer, equilibrated with 1 column volume of PBS pH 7.4, and then incubated with 1 µM His-tagged *Cth* Ulp1 protease for 1–2 hours (*1*) at 25°C with mild shaking to suspend the Ni-NTA resin. The eluent was collected. The resin was rinsed with an additional 1 column volume of PBS pH 7.4, which was collected and pooled with the eluent.

### Preparation of Fluorescein-conjugated Protein

5-(iodoacetamido)fluorescein (5-IAF) was purchased from Abcam, dissolved to 20 mM in DMSO, and stored in aliquots at –20°C until use.

Mutants containing a single cysteine residue were cloned from an eblock as described under *DNA Cloning*. The proteins were expressed and purified as described under *Protein Expression and Purification* with two key modifications. First, during the lysis and washing steps, 1 mM of fresh tris(2-carboxyethyl)phosphine (TCEP) was added to Lysis Buffer I and Wash Buffer to reduce any disulfides. Second, between the washing and elution steps, 5-IAF was conjugated to the free thiol groups via the following steps:

The Ni-NTA magnetic beads were equilibrated into Conjugation Buffer (50 mM Tris-HCl at pH 8.5, 100 mM NaCl, 1 mM TCEP), and the supernatant was removed to leave a small residual volume of buffer with the magnetic beads (still bound to SUMO-tagged proteins). A freshly thawed aliquot of 5-IAF was dissolved 1:50 into Conjugation Buffer to a final concentration of 400 mM, and 200 µL of this solution was added to the magnetic beads. The beads were shaken in the dark at 25°C overnight (16–20 hr), with an appropriate rotation speed to keep the magnetic beads suspended. Afterwards, the supernatant was removed, and the beads were washed 5 times with 2 mL of PBS pH 7.4 to remove excess unreacted dye. From here, the fluorophore-labeled protein was eluted with *Cth* Ulp protease as described under *Protein Expression and Purification*. Single-dye labeling was verified with MALDI-TOF mass spectrometry.

#### Fluorescein Fluorescence Anisotropy Measurement

The UV-vis absorption spectrum of the fluorophore-labeled protein was measured using a NanoDrop (Thermo Fisher), and the concentration was estimated using the absorption at 491 nm and an extinction coefficient of 82,000 M^-1^ cm^-1^ for fluorescein. A standard 1 cm path length cuvette (Agilent 8453 G1103A spectrophotometer) gave absorbance readings within 10% of the value determined on a NanoDrop.

Like most camptothecin derivatives, exatecan contains a lactone ring that hydrolyzes in neutral or basic pH over a timescale of hours (*2*). Binding to EPIC has been previously shown to dramatically slow hydrolysis and stabilize the lactone form (*3*). We conducted all experiments with exatecan or belotecan freshly diluted from a 10 mM DMSO stock (100% lactone form) into PBS at pH 6.5. At pH 6.5, the lactone ring hydrolyzes more slowly than at pH 7.4, and the lactone-carboxylate equilibrium is shifted further towards the lactone form.

To prepare exatecan-bound and exatecan-free protein under otherwise identical conditions (Fig. 2A, 4B, and 4E; tables S4 and S7), we mixed a dilute protein solution 1:1 with either a ligand solution or a buffer with an equivalent amount of DMSO. The final concentrations were 75 nM protein and 5 or 0 µM exatecan in PBS 6.5 with 0.01% w/v PEG-3350 and 0.05% v/v DMSO. The mixture was incubated in the dark at room temperature to allow binding to reach equilibrium. Anisotropy was measured after 30 minutes and after 24 hours.

The fluorescence anisotropy of the lid-attached fluorescein was measured in 384-well plates in a plate reader (BMG LabTech PHERAstar FSX) using an excitation filter of 485 nm and an emission filter of 520 nm. For calibration, we adjusted the relative gains of the parallel and perpendicular emission channels so that a well containing free fluorescein in PBS pH 7.4 had a polarization of 0.035 (anisotropy of 0.024). The measured values of fluorescence anisotropy were constant over a wide range of protein concentrations (1– 1,000 nM). Below ∼1 nM, we found it necessary to subtract off the background values in both channels before calculating anisotropy in order to obtain a concentration-independent value, as described under *Binding Affinity by Fluorescence of Lid-attached Fluorescein*. The same plate reader settings were used for partial proteolysis and binding affinity experiments that measured the fluorescence anisotropy of lid-attached fluorescein.

### Partial Proteolysis Measurement

Lyophilized porcine pancreatic trypsin was purchased from Fischer Scientific (#AAJ6040209) and reconstituted at a concentration of 5 mg/mL in a solution of 1 mM HCl and 20 mM CaCl_2_. Aliquots were flash-frozen in liquid nitrogen and stored at –20°C.

For kinetic measurements of lid cleavage through the fluorescence anisotropy of lid-attached fluorescein (Fig. 2B, fig. S1 and S5, table S4), the protein was first brought to 100 nM in PBS pH 6.5 with and without exatecan (0 or 10 µM), as described under *Fluorescein Fluorescence Anisotropy Measurement*. Next, a Trypsin Solution was prepared by diluting a freshly thawed aliquot of trypsin to 0.5 mg/mL in 1 mM HCl and 20 mM CaCl_2_ and then diluting to 0.2 mg/mL in PBS pH 6.5. The protein solution and Trypsin Solution were placed in adjacent wells on a 384-well plate, and the plate was placed onto the receptacle of the plate reader (BMG LabTech PHERAstar FSX). Proteolysis was initiated by rapidly and thoroughly mixing the equal volumes of the two solutions with a multichannel pipette (final concentrations are 50 nM protein, 0 or 5 µM ligand, 0.1 mg/mL trypsin, and 4 mM Ca^++^). The measurement was initiated immediately after mixing. Because of the time delay involved with retracting the plate receptacle and initializing the detection apparatus on the instrument, the kinetic measurement incurred a dead time of ∼17 seconds. Measurements also incurred a well-dependent time delay because the plate reader scans each well sequentially. The reported data was corrected for the exact duration of the dead time and time delay for each well. The length of the measurement was limited to 90 minutes to minimize baseline drift due to evaporation (roughly 0.5–1.5 µL/hr).

The procedure for fitting rate constants is described in supplementary text S3 and fig. S5. A trypsin concentration of 0.1 mg/mL was used throughout this study. For the designs with fast proteolysis kinetics, we could only determine a lower bound of the initial proteolysis rate at this trypsin concentration (marked with a < in table S4). The initial proteolysis rate of d6.9 was directly proportional to trypsin concentration over a range of 0.001–0.1 mg/mL.

#### Binding Affinity by Intrinsic Ligand Fluorescence

As one way to determine the equilibrium binding affinity between the designed proteins and the ligand (exatecan or belotecan, Fig. 2E), we serially diluted the protein from a starting concentration of 2 µM while maintaining the ligand at a fixed concentration of 10 nM in PBS pH 6.5 with 0.01% w/v PEG-3350. The protein and ligand were incubated at room temperature for a duration ranging from 5 minutes to 72 hours, depending on the on-rate of the design, until the binding reached equilibrium as determined by a converged binding curve. The curves were measured in triplicate in a low-volume 384-well plate (Corning 4514) in a plate reader (BMG LabTech CLARIOstar Plus), with an excitation filter of 360 nm (20 nm bandwidth) and an emission filter of 460 nm (10 nm bandwidth) to detect the parallel and perpendicular emission.

Because the low concentration of exatecan or belotecan (10 nM) gave a weak fluorescence signal, it was necessary to subtract off the background before calculating the anisotropy to obtain a concentration-independent value. The background values of the parallel and perpendicular detection channels were determined by titrating exatecan from 0 nM to 100 nM and taking the y-intercept of a least-squares fit of the two channels separately. With our instrumentation, 10 nM exatecan gave a signal-to-background ratio of ∼5–8 to 1. The anisotropy and intensity were calculated with the formulas

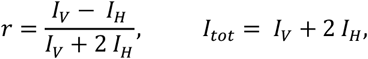

where *I*_*v*_ and *I*_*H*_ are the background-subtracted values in the parallel and perpendicular channels, respectively.

### Binding Affinity by Fluorescence of Lid-attached Fluorescein

For the designs where exatecan binding causes a change in the fluorescence polarization of lid-attached fluorescein (e.g. d6.4 and d6.9), we also determined the binding affinity by fixing the protein concentration and varying the exatecan concentration (Fig. 2D and fig. S4). The brighter emission of fluorescein, compared to exatecan, allowed us to reduce the concentration of the fixed species in the titration curve (0.1 nM fluorescein-conjugated protein compared to 10 nM exatecan with our instrumentation). The labeled protein was kept at a fixed concentration (1.0 nM, 0.33 nM, or 0.11 nM) and exatecan was serially diluted from 100 nM in PBS pH 6.5 with 0.01% w/v PEG-3350. The protein and ligand were incubated at room temperature in the dark for a duration of 30 minutes (d6.4^cpΔ11^), 2 days (d6.9), or 11 days (d6.4 and d6.4^M93E^) in a tightly sealed low-volume 384-well plate (Corning 4514). Within this time frame, designs d6.4^cpΔ11^, d6.9, and d6.4^M93E^ are expected to reach equilibrium (based on the inferred off-rate); however, d6.4 is expected to have an equilibration time of several months, so the measured dissociation curve after 11 days represents an upper bound. Issues of evaporation and/or sample degradation prevented us from extending the incubation time. Fluorescence anisotropy was measured in a BMG LabTech PHERAstar using an excitation filter of 485 nm and an emission filter of 520 nm. Background values in the parallel and perpendicular channels were determined by titrating fluorescein from 1 nM down to 0 nM and taking the y-intercept of a least-squares fit of the two channels separately. With our instrumentation, 1 nM fluorescein gave a signal-to-background ratio of ∼10–15 to 1.

### Fitting K_D_ to Fluorescence Polarization Curves

For binding curves with fixed protein concentration and variable exatecan concentration (Fig. 2D and fig. S4), the binding affinity was fit globally across three protein concentrations (1.0 nM, 0.33 nM, and 0.11 nM) to a single-site binding model that accounts for ligand depletion,

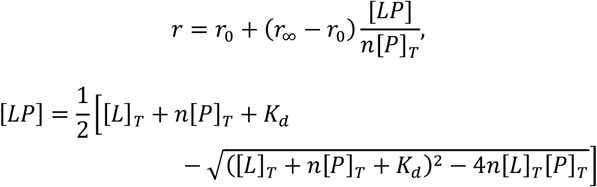

where *r* is the experimentally measured anisotropy, *r*_∞_ is the anisotropy of unbound protein, *r*_0_ is the anisotropy of bound protein, [*L*]_*T*_ is the total ligand concentration added, [*P*]_*T*_ is the total protein concentration added, *K*_*d*_ is the dissociation constant, and *n* is a stoichiometry parameter (used here to account for inaccuracies in protein concentration relative to ligand concentration). The parameter *K*_*d*_ was fitted globally across the three datasets, and the parameters *r*_0_, *r*_∞_, and *n* were fit individually for each dataset with the constraint 0.8 < *n* < 1.2 . We determined the standard error of each fit parameter from the square root of the diagonal entries of the covariance matrix, given by the formula

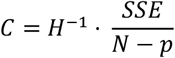

where *SSE* is the weighted sum of squared errors at the optimized parameter values, *N* is the number of data points, *p* is the number of parameters, and *H* is the Hessian of the *SSE* evaluated at the optimized parameter values (based on numerical differentiation).

For binding curves with fixed exatecan concentration and variable protein concentration (Fig. 2E), the binding affinity was fitted with the same equations with the roles of protein and ligand swapped. Only a single ligand concentration was used (10 nM) and we did not perform a global fit. In cases where K_d_ ≲ 2 nM, it becomes difficult to accurately determine K_d_ because the fixed ligand concentration is higher than the K_d_ in the experiments; we reported these values simply as K_d_ < 2 nM.

### On-rate by Intrinsic Ligand Fluorescence

As one way to measure association kinetics of exatecan to our designed proteins, we mixed together the protein and ligand and monitored the subsequent increase in exatecan fluorescence anisotropy on a plate reader (Figs. 2F, 4C, and 4F; fig. S8; table S7). Ligand concentration was held fixed at 25 nM and protein concentration was varied from 0 to 5 µM. A 2x ligand solution and 2x protein solution were prepared in adjacent wells on a 384-well plate, and the plate was placed onto the plate reader receptacle. Next, equal volumes of the two solutions were rapidly and thoroughly mixed with a multichannel pipette, and the kinetic measurement was initiated immediately. The dead time was determined as described under *Partial Proteolysis Measurement*. Kinetic measurements used the same plates, plate reader, and wavelength filter set as the equilibrium binding measurements (see *Binding Affinity by Intrinsic Ligand* Fluorescence).

#### On-rate by Fluorescence of Lid-attached Fluorescein

For the designs where exatecan binding causes a change in the fluorescence polarization of lid-attached fluorescein, we also determined the binding on-rate using anisotropy of the lid-attached fluorescein to track the fraction of protein bound. Protein concentration was held fixed at 10 nM and exatecan was varied from 0 to 40 µM (fig. S8). The experimental procedure follows *On-rate by Intrinsic Ligand Fluorescence*, and the plate reader operation follows *Binding Affinity by Fluorescence of Lid-attached Fluorescein*.

### Fitting on-rates to kinetic association curves

For association kinetics determined by fluorescence of lid-attached fluorescein (fig. S8), the kinetic trace was modeled with the equation

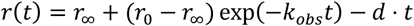

where *r*_∞_ is the fluorescence anisotropy of the lid-attached dye in ligand-free protein, *r*_∞_ is the limiting anisotropy at long time (i.e. the average of free and bound anisotropy weighted by equilibrium fraction bound), *k*_*obs*_ is the observed pseudo-first-order rate constant at a given protein concentration, and *d* accounts for instrument drift over the observation time. This expression assumes that [*L*] ≫ [*P*] so ligand depletion can be neglected (pseudo-first-order condition).

We performed a single-exponential, least-squares fit independently for each of the protein concentrations. Then, we fit the overall on-rate, *k*_*on*_, from the observed rate constants across multiple protein concentrations using the equation

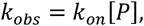

which assumes that [*L*] ≫ *K*_*d*_ so dissociation (i.e. k_off_) can be neglected. The uncertainty was propagated from the first to the second fitting step by weighting each term in the sum of squared residuals.

For association kinetics determined by intrinsic anisotropy of exatecan (Figs. 2F, 4C, and 4F; fig. S8; table S7), the kinetic trace was modeled with the equation

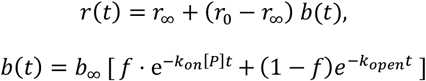

where *k*_*on*_ is the bimolecular rate constant, *k*_*open*_ represents a (slower) concentration-independent lid-opening rate, *f* is the amplitude of the bimolecular component, *b*_∞_ is the equilibrium fraction of bound ligand, and *r*_0_ and *r*_∞_ are the anisotropy values of free and bound exatecan, respectively. The parameters *k*_*on*_, *k*_*open*_, *r*_0_, and *r*_∞_ were fitted globally across protein concentrations, and the parameters *b*_∞_ and *f* were fitted separately for each protein concentration, with constraints 0 < *b*_∞_ < 1 (d6.4^cp^ truncation series) or 0.8 < *b*_∞_ < 1 (d6.4, d6.9, and d6.4^M56E^ based on [*P*] ≫ *K*_*d*_ ). It is expected that two components are necessary to explain the kinetic trace (one [P]-dependent and one [P]-independent) since the overall kinetic scheme involves multiple steps (closed ⟷ open ⟷ bound). The optimal parameter values were determined by minimizing the overall sum of squared residuals across all protein concentrations. We ran the fitting procedure starting from several initializations to escape local minima. Standard errors are determined as described under *Fitting K_d_ to Fluorescence Polarization Curves*.

### Protein Crystallization

A fresh preparation of protein was buffer exchanged into PBS pH 6.5 and concentrated to 230 µM with spin filtration (3K Amicon ultra centrifugal filters; Millipore). The protein was then mixed with a 1.2-fold molar excess of exatecan and allowed to equilibrate for 10 minutes (d5.11) or 1 hour (d6.9 and d6.4). The solution then was concentrated further via spin filtration to a final concentration of 3.5 mM (70 mg/mL). Then, 0.2 µL protein was mixed 1:1 with mother liquor and set as sitting drops in a 96-well Intelli-plate (Hampton Research HR3-185) using an NT-8 Robot (Formulatrix). Crystallization conditions were tested from the sparse matrix screens MCSG-2 and MCSG-3. The conditions that produced crystal growth are listed in table S5. Crystals were looped and dipped briefly in a cryoprotectant of Paratone-N before plunging into liquid nitrogen.

### X-Ray Diffraction and Structure Refinement

X-ray diffraction datasets were collected at 100 K at beamline AMX at NSLS-II (Brookhaven National Lab). The data were indexed, merged, and scaled using autoPROC (*4*) or fast_dp (*5*), and the structures were phased by molecular replacement with PHENIX (*6*) using the coordinates from the AlphaFold3 prediction. Multiple rounds of model building and refinement were performed using Coot (*7*) and PHENIX. Resulting data collection and refinement statistics are summarized in table S6.

### FRET emission spectra

Designed protein fused with fluorescent proteins, mTurquoise2 and mVenus, was expressed and purified as described under *Protein Expression and Purification*. The concentration was determined using the absorption at 515 nm and an extinction coefficient of 104,000 M^-1^ cm^-1^ for the mVenus chromophore. SDS-PAGE gels showed no cleavage between protein domains. The protein and ligand were prepared at 2× the final concentration, mixed in a 1:1 volume ratio, and equilibrated at room temperature for 30 min before measurement. For measurements in saliva, urine, and serum, the protein was prepared in PBS pH 6.5 with 0.01% w/v PEG-3350, and the ligand spiked into the biological fluid. Saliva and urine samples were obtained from a healthy human donor and briefly centrifuged before use. Human serum was purchased from Millipore Sigma (H3667).

The emission spectra were measured in 384-well plates in a BMG LabTech CLARIOstar plate reader, which uses a diffraction grating to select excitation and emission wavelengths. We used an excitation wavelength of 431 nm with a bandwidth of 8 nm, and we detected the emission over the spectral range 450–580 nm with a bandwidth of 8 nm and a stepwidth of 1 nm. We processed the raw spectra by subtracting off the background from environment and applying a smoothing window of 3 nm. The donor-to-acceptor peak ratio was calculated from the resulting smoothed spectra.

The binding affinity of mT-d6.4^cpΔ11^-mV to exatecan was determined through a global fit across three protein concentrations (32 nM, 10 nM, 3.2 nM) using the FRET ratio to track the fraction of protein bound (fig. S12). The parameters *K*_*d*_, *r*_∞_, and *r*_0_ were fitted globally across the three datasets, and a separate parameter *n* was fitted individually for each dataset with the constraint 0.8 < *n* < 1.2, as described under *Fitting K*_*d*_ *to Fluorescence Polarization Curves*. The titration curves in Fig. 4B are fit to the same equation using the K_d_ determined from the global fit (1.5 nM) and separately fit values for *r*_∞_, *r*_0_, and *n* in each condition (with the constraint 0.9 < *n* < 1.1 ). For binding curves in saliva, urine, and serum, we included an additional parameter [*A*]_*T*_, the concentration of environmental proteins that binds competitively to exatecan. The binding curve is fitted to a numerical solution to the system of equations

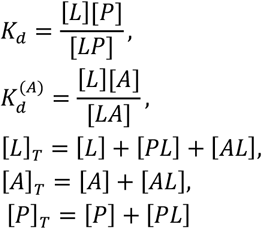

where we fixed *K*^(6)^ to 43 µM, the affinity of human serum albumin (HSA) to exatecan (*3*). In saliva or urine, where HSA may be found in low concentrations (or not at all), we still used a *K*_d_ of 43 µM to model weak, non-specific binding of the drug to matrix proteins.

## Supplementary Text

### S1. Computational Design Methods

#### Seeding lids using Query-Based Interfacial Tertiary Structures (QBITS)

Our design process started with the 4-helix bundle, EPIC (pdb 9NZE), which binds exatecan in the bundle interior. [Except in reference to this crystal structure (9ZNE), throughout this paper, EPIC refers to the (M93L, Q51N) double mutant with increased affinity (K_d_ = 1.2 nM).] We sought to design a lid domain that interacts with both the solvent-exposed portion of exatecan as well as the protein surface. To do this, we developed a structural bioinformatics algorithm called QBITS (query-based interfacial tertiary structures), which uses a structure-based search of a protein database to rank prevalent interfaces that pack against a structural query.

We first created a non-redundant database of the protein data bank (PDB), using biological assemblies instead of single chains to maximize the number of interfaces in the database. The QBITS database is 20,567 protein crystal structures from the PDB (before Aug 2020), filtered by resolution < 2.8 Å, R_free_ < 0.35, and sequence identity < 70% across biological assemblies (after concatenating sequences across chains and considering permutations). When parsing through search results, further redundancy in structural matches is handled internally by the QBITS software (to collapse identical interfaces from the same biological assembly for frequency statistics).

The QBITS software parses through the results of a structural search through this database from the program MASTER (*8*). Matches to a structural query that have a Cα RMSD less than a user-defined threshold (here, 1.2 Å) are tabulated by MASTER, including the matched residue indices in the PDB file, the sequence of the matched residues, and the RMSD of the match. QBITS takes this information, and for each match, superposes the matched protein residues to the query. In our case, the query was two helical segments that sandwich exatecan (12 residues on α4 and 12 residues on α1, fig. S7 and S14), resulting in 11,122 unique matches. QBITS then removes any residues of the superposed proteins that clash with the entire query protein (in our case, EPIC) or that are further than 20 Å from any residue in the query.

Next, QBITS breaks the remaining interfacial residues from each match into contiguous sliding windows of a user-defined length (here, 7 residues). Any contiguous fragments with fewer residues are dropped. For lids to EPIC, we also dropped any fragments that were not helical (DSSP assignment of H for each residue). The Cα coordinates of the remaining residue windows, across all proteins with structural matches to the query, are stored in a ball-tree data structure that supports rapid nearest-neighbor queries. Using this tree, QBITS scores each residue window by the number of nearest neighbors within a Cα RMSD threshold (here, 1.5 Å).

Next, QBITS nominates candidate interfaces from the matched proteins as collections of residues with high sliding-window scores. A QBITS interface can be a single contiguous segment or a concatenation of discontiguous segments (with minimum length of a single sliding window, here 7 residues). Each QBITS interface is given an aggregate score of Σ_i_ log(*N*^*(i)*^_nbrs_ + 1), where *N*^*(i)*^_nbrs_ is the number of nearest neighbors of window *i* and the sum runs over all contiguous windows contained in the interface for a single matched PDB file. Interfaces are ranked by this score (higher is better), and the highest scoring interfaces are chosen as QBITS “representatives”. This score biases candidate interfaces toward those with more neighbors (higher prevalence in the PDB) and more contiguous residue windows (larger interfaces). Here, we limited QBITS output to single contiguous segments of arbitrary length, with the additional criterion that each 7-residue sliding window within the segment needed to have at least 3 nearest neighbors. We used these contiguous representatives (250 total) from QBITS as initial single-helix seeds for the lids to EPIC (leading directly to designs d5.1-d5.12). In the design of 2-helix lids (see below), we also used contiguous QBITS representatives but additionally used low-RMSD clusters of sliding windows (derived from greedy clustering of the ball tree with a Cα RMSD cutoff, e.g. 1 Å) to more finely sample helix coordinates centered around top QBITS representatives.

Overall, we found QBITS to be a rapid and straightforward way to compute potential interfaces for subsequent design. A QBITS computation, together with the initial MASTER search to identify structural matches, takes about 5 min on a MacBook M1 CPU to generate potentially thousands of ranked interfaces for a given query. Compared with RFdiffusion, QBITS is a computationally inexpensive method to rapidly generate interfaces based on known examples from the PDB; it does not rely on slow, iterative denoising to generate coordinates using GPUs. Because QBITS interfaces are derived directly from crystal structures, the data provenance is easy to track, unlike from a generative neural network. Lastly, filtering for helical content and directionality is straightforward, producing output that satisfies user-defined parameters.

#### Filtering seeds for potential salt bridge using van der Mers

Next, we filtered the list of 250 helix placements based on whether it was possible for one of its residues to harbor a Glu or Asp to form a salt bridge with the exatecan amine. To determine whether such an interaction was geometrically feasible, we used van der Mer (vdM) databases of Glu or Asp residues with the amine chemical group (derived from Lys side chain) (9). Each vdM represents an observed interaction geometry between an amine and the N-C_α_-C backbone frame of a Glu (or Asp) residue. We took the subset of these vdMs that met two criteria: (i) the φ/ψ angles of residues *i-1,i*, and *i+1* with respect to the Glu (or Asp) residue corresponded to an α-helix, as determined by DSSP (*10*); (ii) the -NH_3_ ^+^ group was situated within hydrogen-bonding distance of the -COO^-^ group of the Glu (or Asp) side chain, as determined by the program Probe (*11*). After filtering by the φ/ψ and H-bonding criteria, there were 4,683 (or 3,439) remaining vdMs in the dataset.

For each candidate helix placement from QBITS, we iterated through each residue and determined whether its backbone frame was geometrically compatible with a salt bridge to the exatecan -NH_3_ ^+^ group. To do this, we applied a rigid-body transform to every vdM in the dataset to superpose the N-C_α_-C backbone frame onto each residue in the helix, and we compared the transformed coordinates of the -NH_3_ ^+^ group with the corresponding atoms of exatecan in EPIC. A vdM was considered a structural match if, after superposition, it had a RMSD of < 0.6 Å between the C_γ_, C_δ_, and N_ε_ atoms in the vdM (the amine atoms) and either the C6, C5, and N2 or the C4, C5, and N2 atoms of exatecan (fig. S14). For practical efficiency, we queried the entire vdM dataset at once by first transforming all backbone frames to a common reference frame, storing the -NH_3_ ^+^ coordinates in a ball tree. We accepted a seeded helix from QBITS if at least one of its residues matched with at least one vdM. In total, 58 out of 250 placements passed this filter, including the most prevalent seeds.

#### Filtering for helical crossing angle

We next used a geometric filter of the QBITS representatives to ensure that the seeded helix was oriented along the same axis as the two antiparallel helices of EPIC (α1 and α4) that sandwich exatecan (see fig. S7A for helix labels and fig. S14). Such an orientation ensures that the lid helix can be extended, without excessive strain, to form a larger interface with the binding domain (see next step below). To calculate the crossing angle cosine between two helices, we fitted a helical axis vector to the C_α_ coordinates and took the dot product between normalized axis vectors. We accepted candidate helical placements that formed a crossing angle of < 37° with the query fragment of either α1 or α4. In total, 45 out of 58 seeds from QBITS passed this filter, ranging in length from 9–16 residues.

#### Helical extension with RFdiffusion All-Atom

To form a larger interface between the seeded lid and the binding domain, we extended the length of the lid helix using RFdiffusion All-Atom (RFAA) (*12*). The helix was extended towards the direction of its eventual attachment point on the end of the 4-helix-bundle binding domain. To determine the number of residues to add, we calculated the C_α_ distance (in Å) between the attachment point (terminus of the 4-helix-bundle domain) and the terminus of the lid helix; we converted this length into a residue count, *n*, by dividing by the rise of the α-helix (1.5 Å per residue). The number of residues added to the lid was sampled uniformly from *n* − 3 to *n* + 5 via the contig string input parameter in RFAA. Depending on whether the lid helix was parallel or anti-parallel to α1, the additional residues were added either C- or N-terminally to the lid, respectively.

A total of 12 lid extensions were sampled per QBITS seed. Most of the additional residues generated by RFAA were helical, but occasionally a few residues near the end were non-helical; non-helical portions (defined as a DSSP annotation other than H, I, or T) were removed from the backbone template. The extension process resulted in a total of 540 candidate lids ranging in length from 9 to 29 residues.

#### Sequence design with LASErMPNN

Next, we used the graph neural network LASErMPNN (*13*) to design an amino-acid sequence to support the necessary interactions between the lid, the ligand, and the binding domain.

Within the binder domain, we redesigned a total of 23 positions of EPIC at the protein surface near the interface with the lid. Most residues directly contacting exatecan in EPIC were held fixed, with the exception of E132, a surface residue on α4 which forms a salt bridge to exatecan’s amine (Fig. 1C, upper inset). We sought to allow LASErMPNN to transfer this salt-bridge interaction onto the lid to strengthen the lid–ligand interaction energy. At this stage, we also introduced the mutations Q51N and M97L of EPIC, which were previously found to improve the affinity to exatecan (K_d_ from 120 nM to 1.2 nM) in the original 4-helix bundle (*13*).

In addition to unconstrained sequence design by LASErMPNN (of all lid residues and 23 surface residues of the binder domain), we undertook a parallel effort of explicitly designing a lid–ligand salt bridge via Glu or Asp / amine vdMs. The Glu/Asp side chain with the lowest amine RMSD was grafted onto the structural template, and the modeled side-chain rotamer was provided as a fixed residue in the input structure to LASErMPNN. The remainder of the lid (along with the 23 designable surface residues of the binder domain) was designed by LASErMPNN. We included both the unconstrained set of sequences and the fixed-vdM set of sequences in the subsequent filtering steps.

#### Multi-state design

To further increase sequence diversity, we also sampled sequences using a multi-state design strategy. We took a weighted average of the amino-acid probabilities between a lid-closed, exatecan-bound model and a lid-open exatecan-free model. To create a design model of the drug-free protein with an open lid, we translated the lid helix 20 Å from the binding domain. By separating α5 from α1–4, we sought to encourage LASErMPNN to design a soluble lid that down-weights hydrophobic residues at the binder–lid interface. We note, however, that it is biophysically implausible for the lid to adopt a helical structure without packing interactions with the binder domain (*14*); furthermore, because an isolated α-helix is rarely seen in the PDB, the structural model was outside the training distribution of LASErMPNN, causing it to occasionally generate hydrophobic (e.g., all-Leu) lid sequences. Nevertheless, this multi-state strategy increased sequence diversity, and we included the sequences generated by this weighted average procedure in subsequent filtering steps.

In total, we sampled 80 sequences per lid placement: 4 choices of weighting between the lid-open, exatecan-free and lid-closed, exatecan-bound structures (0.0, 0.5, 0.8, 1.0); 2 choices of sequence constraints (fixing salt bridge or not); and 10 samples per parameter set. We used a sampling temperature of 0.25 and the model weights of LASErMPNN that were trained using the subset of the PDB employed in the LigandMPNN training split (model weights file `ligandmpnn_split_last_chance_edge_vecs_optstep_5 5000.pt`)

#### AlphaFold3 prediction of structures

To rank and evaluate the 43,200 candidate sequences for single-helix lid designs, we used AlphaFold3 to predict the structure of each sequence with and without exatecan bound. A flexible linker of (GGS)_7_ was inserted between the lid and binder sequence to create single-chain 5-helix-bundle sequences. AlphaFold3 was run in single-sequence mode (no multiple sequence alignment) using 5 samples from 1 seed and the following SMILES string for protonated exatecan:

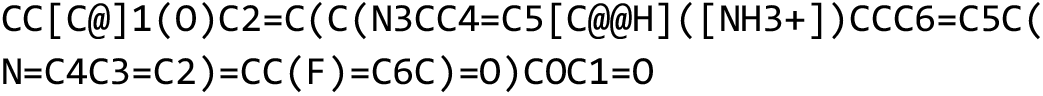

Out of 5 predicted models, only the one with the highest AlphaFold3 `ranking_score` was analyzed. The predicted structure was first aligned to the design model by the Cα coordinates of the binder domain (RMSD < 1 Å for all sequences). In this reference frame, we calculated the RMSD between the predicted lid structure and the coordinates of the closed lid in the design model (i.e., the RFdiffusion extension of the QBITS seed). For each predicted structure (exatecan-bound and exatecan-free), we also calculated the average interfacial predicted aligned error (iPAE) between all pairs of binder and lid residues (Fig. 1D).

We calculated the hydrophobicity of the interface based on the output of LASErMPNN, which, in addition to designing the protein’s sequence, also predicts side-chain rotamers. We used freesasa (*15*) to calculate the interfacial area between the lid and binder domains using the formula

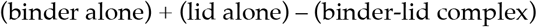

and then calculated hydrophobicity by dividing the apolar interfacial area by the total interfacial area.

#### Filtering of 5-helix designs

To filter for designs that were closed with exatecan bound and open without, we applied the following filtering cutoffs:

- iPAE with exatecan < 5
- iPAE without exatecan > 15
- lid RMSD (AlphaFold3 vs design) with exatecan < 3
- lid RMSD without exatecan > 3

A total of 25 sequences passed all 4 filters. We re-predicted each of the 25 sequences with varying number of GGS repeats in the flexible linker ((GGS)_n_ with n = 3–7) and chose the linker length that maximized the difference in iPAE between the exatecan-bound and exatecan-free predictions. We selected 12 of these 25 proteins, named d5.1–d5.12, based on robustness to linker length, overall interfacial size, similarity to other designs, and prediction of no homo-dimerization by AlphaFold3 (high PAE between two identical chains).

#### Design of 2-helix lids

The design of 2-helix lids (6-helix proteins), d6.1–d6.12, followed a similar computational protocol as for single-helix lids (5-helix proteins). Initial experiments with d5.1–d5.12 identified d5.11 as a weak hit by the limited proteolysis assay (fig. S1) but not by equilibrium fluorescence anisotropy. The small magnitude of the ligand-dependent change in protection from cleavage by trypsin suggested that the lid interacted with the ligand too weakly to lead to any substantial conformational shift detectable by equilibrium anisotropy. We solved a crystal structure of exatecan-bound d5.11 (fig. S6), which showed a portion of the ligand surface remained solvent-exposed (10 Å^2^ in d5.11 compared to 50 Å^2^ in EPIC, calculated using freesasa). We reasoned that adding a sixth α-helix (called α6) would further bury the ligand and increase the lid–ligand interaction strength (see also supplementary text S4).

The design of 2-helix lids proceeded in two phases: an initial broad search with QBITS to identify promising α6 placements, and a more targeted QBITS-based search to focus on the search space near the top α6 placements. The two phases are labeled I and II in Table S3.

#### Placement of α6 helix

We found candidate seeds of the α6 helix using the same QBITS-based workflow as the α5 helix (see fig. S7 for helix numbering scheme). As the structural template, we used the exatecan-bound crystal structure of d5.11. The structural search query for MASTER was comprised of two helical segments (11 residues of α5 and 12 residues of α1, fig. S14). Using the same query RMSD cutoff (1.2 Å) as used previously, we found a total of 4,786 unique structural matches. We filtered the resulting interfaces to those with helical segments parallel to α1. We did not apply the vdM-based geometric filter. Following the same helix scoring procedure, we chose the top 45 contiguous QBITS representatives as α6 helix placements, which ranged in length from 7 to 16 residues.

#### α5–α6 loop design with RFdiffusion All-Atom

In contrast to the flexible (GGS)_n_ linker between α4 and α5, we designed a structured loop connecting α5 and α6 to form a cohesive lid domain. We used MASTER to determine optimal loop lengths by searching for structural matches of 7-residue segments of paired α5 and α6 in a single-chain non-redundant database of the PDB (RMSD cutoff of 0.8 Å). We sampled loop lengths in RFdiffusion All-Atom using the gap-length distribution between the matched helix pairs from MASTER. This generated up to 40 loop backbones for each of the α6 placements. After joining the α6 and α5 helices, we had a total of 1,592 structural templates for sequence design.

#### Sequence design of α6

We used LASErMPNN to design the interactions between α6 and the rest of the protein (weights `laser_weights_0p1A_noise_ligandmpnn_split.pt`). In addition to α6 itself, along with the loop connecting α6 to α5, we allowed residues in close spatial proximity to α6 to be designable, localized primarily to α1 and α5. We sampled 10 sequences for each structural template, giving a total of 15,920 sequences.

#### AlphaFold3 filtering

To rank the designed 6-helix-bundle sequences, we used AlphaFold3 to predict the structure of the exatecan-bound complex. We aligned the predicted structures to their corresponding design models by superposing α1–4 (C_α_ RMSD within 1 Å for all designs).

Our first two filtering metrics were (1) a low C_α_ RMSD of α5 and (2) a low average pAE between α5 and α1–4. We elected to use the same cutoffs as the 5-helix designs (RMSD < 3 and pAE < 5). Most candidate designs (12,981/15,920) passed these filters (table S2, column I). Next, we filtered remaining designs for (1) C_α_ RMSD of α6 < 4 Å and (2) average pAE between α6 and α1–4 < 10. After this filtering step, 3,764 of 12,981 sequences remained. We then used the AlphaFold3 predictions to evaluate the hydrogen-bonding geometry between the Glu on α5 and exatecan’s amine. We used two geometric criteria: (1) a distance of < 3.5 Å between the Glu C_δ_ in the AlphaFold3 prediction and the exatecan N2 in the superposed (via α1–4) design model (AlphaFold3 slightly moves the ligand ∼ 1 Å relative to its location in the crystal, fig. S6), and (2) a distance of < 1.5 Å between the Glu C_α_ in the design model and the Glu C_α_ in the predicted structure. Only 24 of 3,764 sequences passed these filters. As a final filter, we inspected designs for packing around the Glu sidechain that permitted no alternate rotamers, resulting in 7 sequences nominated for experimental testing.

#### Sampling more thoroughly around top solutions

We observed that all of the 7 chosen 6-helix sequences shared the following characteristics: (1) they were derived from only 2 of the original 45 seed α6 placements (which QBITS ranked as #1 and #8); (2) the α5–α6 loop length was 9 or more residues in each design; (3) each selected design featured a helical extension of the α5 helix by 3 or 4 residues (from the RFdiffusion step). To more thoroughly sample the nearby structural space, we repeated the previously described computational workflow with the following two modifications:

1. In the α6 placement step by QBITS, we sampled structural diversity around QBITS #1 and #8. We ran a greedy algorithm to cluster all 7-residue contiguous interface windows in the QBITS nearest-neighbors ball tree (RMSD cutoff 1 Å here), and sampled additional α6 seeds from the two clusters overlapping with QBITS representatives #1 and #8 (cluster sizes 16 and 8, respectively). In total, we included 24 new seed placements of the α6 helix.
2. In the α5–α6 loop generation step, we changed the loop length from 0–15 residues to 9–25 residues. We reasoned that a longer insertion length would make it more likely for RFdiffusion to generate an extended α5 helix.

The filtering statistics for the second search phase are summarized in column II of Table S3. We selected 5 of the final sequences from search phase II for experimental testing. Across the two search phases, we chose a total of 12 sequences, d6.1–d6.12, for experimental testing.

### S2. Discussion of Fluorescence Anisotropy as an assay for lid closure

The fluorescence anisotropy of a lid-attached probe reflects the magnitude of local, nanosecond-timescale fluctuations (*16*). In our initial screen of 24 designs, the wide range of anisotropy (0.094–0.239) reflects variation both in the degree of lid closure and in the immediate fluorophore environment. While high probe anisotropy is a strong indication of a rigidly structured lid, low probe anisotropy can be attributed to increased conformational flexibility without complete loss of secondary structure. Furthermore, low probe anisotropy does not rule out the possibility that the fluctuations responsible for depolarization are localized to a segment of the lid near the probe. A holistic picture of the conformational ensemble of the lid requires additional biophysical techniques, such as limited proteolysis, which reports on unfolding events (supplementary text S3), or on-rate measurements, which reports on lid-opening motions that are large enough to allow ligand entry.

### S3. Discussion of Limited Proteolysis as an assay for lid closure

Limited proteolysis provides a convenient method to assess ligand-induced lid closure. Compared to equilibrium fluorescence anisotropy, limited proteolysis is more sensitive because it allows the open state to be detected even if it comprises only a small subpopulation of the conformational ensemble (e.g., as seen with d5.11). Under certain assumptions, the rates of proteolysis can also be used to infer the energetics of the lid–ligand interaction, as discussed below.

*Background*. When a polypeptide chain is exposed to a nonspecific protease such as trypsin, proteolysis occurs rapidly in regions that are loosely structured and easily accessible. By contrast, tightly folded regions are resistant to proteolysis unless a conformational fluctuation opens the polypeptide chain to expose a potential cleavage site. In many proteins, binding of a ligand causes a region of the protein to become more tightly structured and thereby more resistant to proteolysis. For instance, in the enzyme iodotyrosine deiodinase, substrate binding causes a lid to close over the active site, leading to a 160-fold reduction in the rate of proteolysis of the lid (*17*).

In our lidded designs, the covalent fluorescein dye allowed us to monitor the kinetics of proteolysis in real time (*18*). Cleavage leads to a sharp drop in fluorescence anisotropy because the resulting dye-attached fragment has a much smaller size than the original protein. The advantage of this fluorescence-based readout compared to standard gel electrophoresis was that it permitted higher-throughput testing of several designs on a plate reader. We attributed decay in anisotropy over time to cleavage of the lid because the (GGS)_n_ linker contains no tryptic sites (Arg or Lys) and the binder is an extremely thermostable de novo designed 4-helical bundle (the parent protein, EPIC, had a T_m_ > 95°C (*13*)).

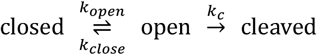

with the assumptions that there is only one cleavage step and only one protease-accessible “open” state. The parameter *k*_c_ describes the intrinsic cleavage rate of a fully-open lid (the 4-helix bundle itself is entirely resistant to proteolysis under these conditions) and depends on the protease concentration and the number of cleavage sites on the lid. Based on this kinetic scheme, the initial rate of cleavage is

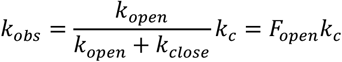

where *F*_*open*_ is the equilibrium fraction in the open state, which is related to the equilibrium constant as *F*_*open*_ = *K*_*open*_/(1 + *K*_*open*_). If the parameter *k*_*c*_ can be measured (e.g. from the proteolysis rate of a model fully-open substrate), then the open fraction *F*_*u*_ can be inferred from the rate ratio *k*_*obs*_ /*k*_*c*_.

Depending on whether the rate-limiting step is opening or cleavage, the overall reaction rate can reflect either the kinetics of lid opening or the thermodynamics of the open/close equilibrium (analogous to EX1 and EX2 in hydrogen-deuterium exchange). The two limiting regimes have a different steady-state rate but the same initial rate, *k*_*obs*_. To avoid complications in interpretation, we chose to fit the initial rate instead of the entire kinetic trace. Focusing on the initial rate also allowed us to avoid analyzing multiple, sequential cleavage events.

Suppose that the binding of the ligand stabilizes the closed state over the open state by a free energy difference of ΔΔ*G*_*lig*_ (defined in fig. S4). The parameter ΔΔ*G*_*lig*_ represents the energy of interactions between the lid and the ligand. Due to this lid–ligand interaction energy, ligand binding will shift the open/close equilibrium away from the open state and reduce the proteolytic rate. Thus, to the extent that this simple scheme is valid, the ligand-induced difference in proteolytic rate can be used to infer ΔΔ*G*_*lig*_.

#### Interpretation of proteolytic kinetics of d6.4 and d6.9

Fig. S4D shows the kinetic traces for the proteolysis of d6.4 and d6.9 with and without exatecan bound. In both proteins, the binding of exatecan clearly protects the lid from proteolysis. We quantified the initial rate of proteolysis by dividing the initial slope by the overall change in anisotropy (initial – cleaved). Using the thermodynamic relations based on this simple kinetic model (fig. S4C), we inferred ΔΔ*G*_*lig*_ of 2–3 kcal/mol for the two designs.

We also measured the proteolytic rate of fully-open control substrates, d6.4^EPIC^ and d6.9^EPIC^, under the same experimental conditions. Because these control proteins share the same amino-acid sequence of the lid as the lidded designs, the tryptic sites are identical and the rate of the control protein can be compared directly to the lidded design. Since the binding domain in the control protein has been replaced with EPIC, the closed state is inaccessible and the lid can be assumed to adopt a purely open conformation.

As expected, d6.4^EPIC^ and d6.9^EPIC^ are proteolyzed more rapidly than d6.4 and d6.9 (fig. S4D). Under the kinetic model, the ratio of initial rates between the lidded design and the control protein corresponds to the equilibrium fraction in the open state. Comparing d6.4 and d6.9, we observed that the proteolytic trace of d6.9 is rather close to d6.9^EPIC^, whereas for d6.4 the proteolysis proceeds notably slower than its fully-open control (fig. S4D). This observation indicates that d6.9 is more open than d6.4. This difference between d6.4 and d6.9 was observed not only in the limited proteolysis assay; in both the fluorescence anisotropy of a lid-attached dye (at t = 0 in this assay) and the rate of association to exatecan (Fig. 2F), d6.9 behaves more similarly to its fully-open control than does d6.4.

#### Screening designs with limited proteolysis

The proteolytic traces of all 24 designs, with and without exatecan bound, are plotted in fig. S1 and summarized in table S4. Tryptic cleavage rates vary widely across the 24 designs, reflecting variation both in the conformation of the lid and in the number of cleavage sites. For 9 of 24 designs, binding of exatecan causes a significant reduction in the lid cleavage rate. Our kinetic scheme relates the ligand-dependent change to a lid–ligand interaction energy, but even at a qualitative level, it is clear that the bound ligand stabilizes the lid in a conformation that is more resistant to cleavage. (We observed the same ligand-dependent protection with both trypsin and chymotrypsin; absolute rates differed but the relative ligand-dependent ratio was comparable.)

#### Interpretation of the protease-susceptible state

Proteolysis is sensitive to a different timescale of motions than the nanosecond-timescale fluctuations probed by a covalently attached fluorescent dye (supplementary text S2). For the lid to be cleaved by trypsin, the polypeptide chain must adopt a conformation that is extended enough to enter the active site of the protease. Such an event requires a large fluctuation from an *α*-helical structure. But the protease-accessible subpopulation might correspond to a complete unfolding into a random coil or to a partial fraying of the end of an *α*-helix. Ultimately, the proteolytic rate reflects the fraction of molecules with any conformation of the lid that is locally unstructured enough to be cleaved by trypsin.

In this work, we found that different experimental techniques, such as equilibrium fluorescence anisotropy and limited proteolysis, largely corroborated each other and provided complementary evidence for ligand-induced lid closure. Indeed, the assays agreed well enough that optimizing the truncation length of d6.4^cp^ via fluorescence anisotropy (to create d6.4^cpΔ11^) led to creation of a construct with a ligand-dependent change in FRET, a completely orthogonal readout.

### S4. Energetic requirements for ligand-induced conformational change

Ligand-induced lid closure can be described with the thermodynamic cycle,

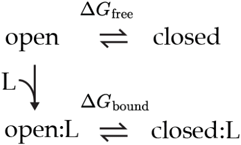

For ligand binding to induce a conformational shift, energetics must be tuned such that both ΔG_unbound_ > 0 and ΔG_bound_ < 0. In addition, the coupling energy, ΔΔG_lig_ = ΔG_bound_ –ΔG_unbound_, must be large enough to result in a substantial population difference (e.g. RT ln(100) = 2.8 kcal/mol for a 100-fold conformational shift). These energetic requirements are general to any design task involving ligand-induced conformational change. In the case of lid closure, the design rationale is particularly straightforward because the two key energetic parameters have clear and distinct structural interpretations.

For lid closure, ΔΔG_lig_ represents the free energy of the interactions between the ligand and the closed lid. The small solvent-exposed surface area of exatecan in EPIC (50 Å^2^) makes it especially challenging to achieve sufficiently strong lid–ligand interactions. We found it was necessary to make complete use of the available ligand surface. In d5.11, where the lid only partially buries exatecan and leaves 10 Å^2^ exposed (fig. S6), ligand binding has only a slight effect on proteolytic resistance (2.6-fold reduction in rate, corresponding to 0.6 kcal/mol, fig. S1). By contrast, in d6.4 and d6.9, where the lid completely buries exatecan, limited proteolysis revealed a much more substantial ΔΔG_lig_ of 2–3 kcal/mol. The crystal structures of d6.4 and d6.9 showed that the lid–ligand interaction energy comes from hydrophobic contacts with Thr44 and a buried salt bridge with Glu41, along with a water-mediated hydrogen-bond between Tyr37 and a carbonyl of the ligand in d6.4.

The other energetic parameter, ΔG_unbound_, is related to the strength of the interface between the lid and the binder. A ligand-dependent conformational shift requires that the interfacial strength be tuned appropriately. If the interface is too strong, then the lid is always closed irrespective of ligand binding (and indeed could preclude binding); if the interface is too weak then the lid is always open. Our experiments showed that this parameter is easily adjustable by truncating the lid or introducing polar substitutions to destabilize the interface, ideally remote from the ligand to avoid disturbing ΔΔG_lig_. The straightforward tunability of ΔG_unbound_ means that weaker hits such as d6.4 can be readily optimized as long as the initial design has a large enough magnitude of ΔΔG_lig_, which can be assessed by limited proteolysis.

## Supplementary Figures

**Fig. S1.**
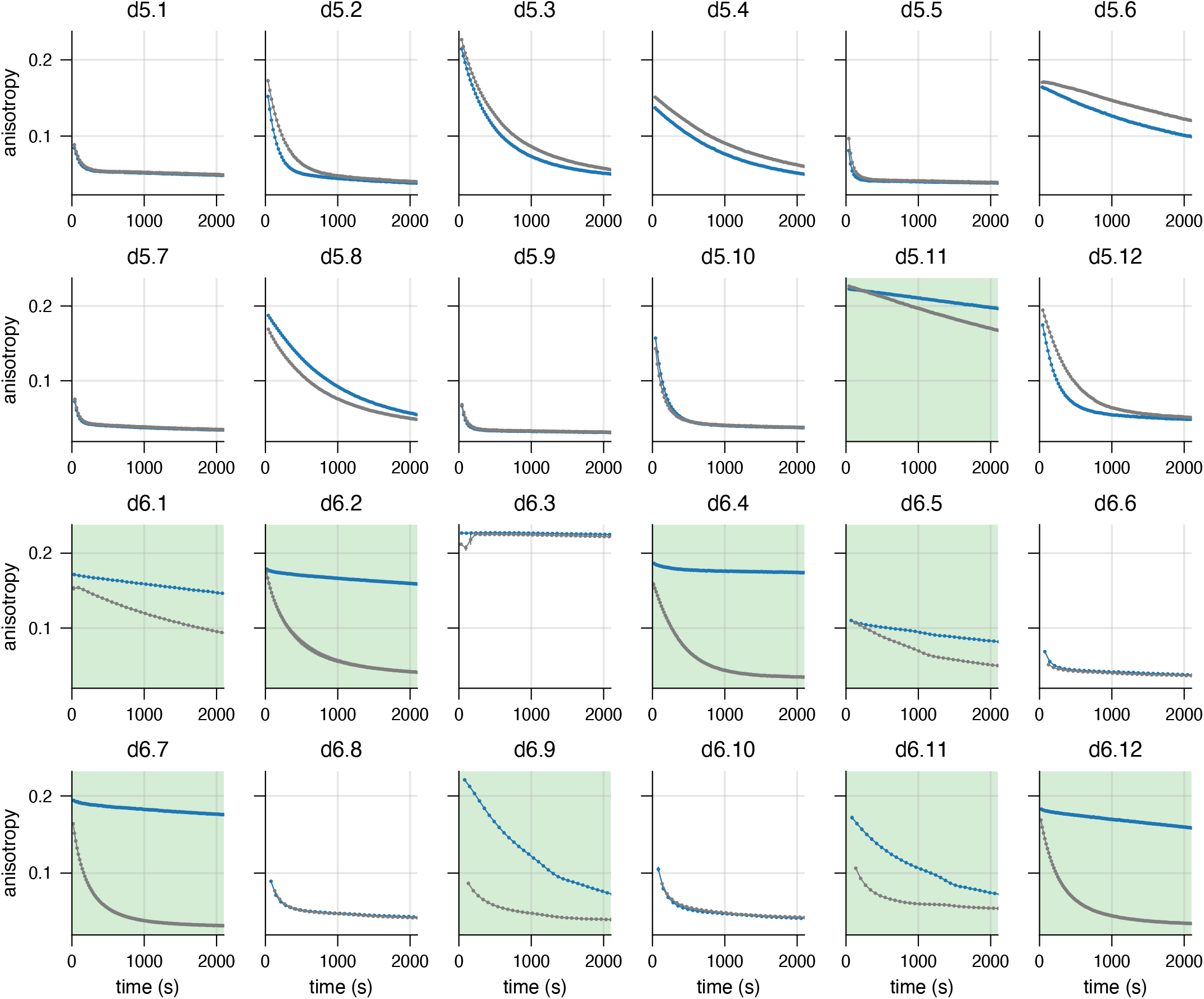
Kinetic traces of limited proteolysis for all 24 designs. As an initial screen for exatecan-induced lid closure, fluorescein-labeled lidded protein (50 nM) is treated with trypsin (0.1 mg/mL), and cleavage of the lid is monitored through the subsequent decay in the anisotropy of the dye label. Each design is proteolyzed in the presence (5 µM, blue curve, saturating ligand) and absence (0 µM, gray curve) of exatecan. The 9 designs with a >2-fold difference in initial rate are colored with a green background.

**Fig. S2.**
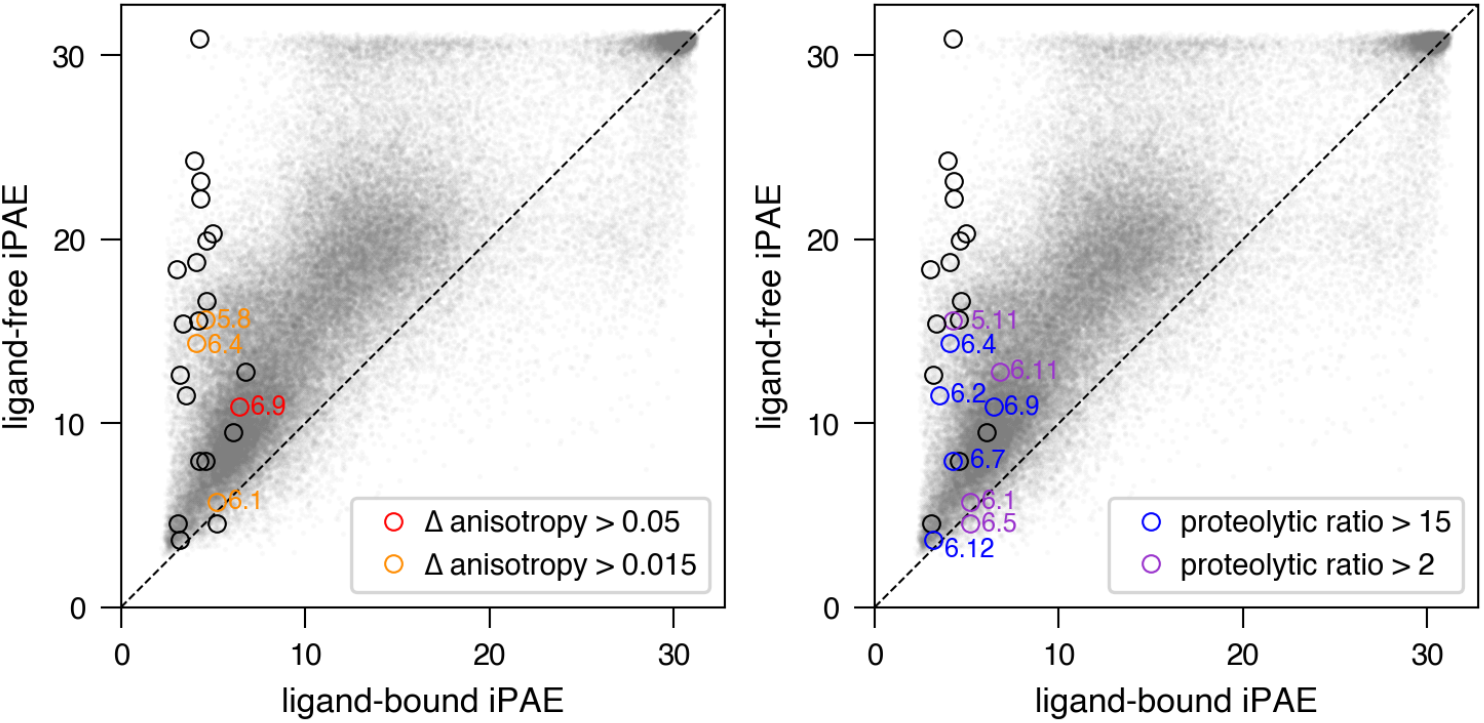
Computational metrics for experimental design hits. The gray scatterplot represents the distribution of computational metrics for all 68,720 candidate designs, with the AlphaFold3-predicted ligand-bound iPAE *vs* ligand-free iPAE. Circles indicate the 24 designs that underwent experimental testing. Colored circles identify the designs that exhibited exatecan-induced lid closure by the equilibrium fluorescence polarization assay (left) or by the limited proteolysis assay (right).

**Fig. S3.**
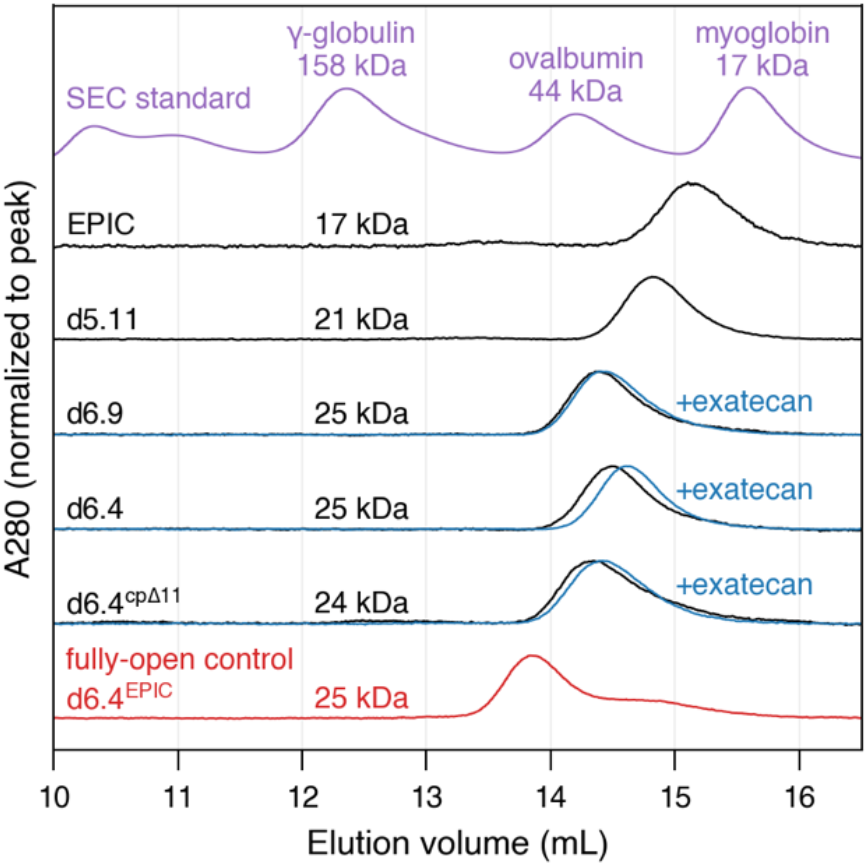
Size-Exclusion Chromatography. Size-exclusion chromatography traces of 2 µM of the indicated proteins using an Enrich 650 column. Each of the indicated proteins is monodisperse. Addition of 10 µM exatecan (blue) to lidded proteins causes a slight rightwards shift in elution volume, consistent with a more compact shape of the protein. Elution volume depends both on the mass and shape of the protein; the oblong shape of the designed helical bundles causes them to pass through the resin more quickly than a more spherical protein of the same molecular weight (e.g., EPIC vs myoglobin). Likewise, the fully-open control d6.4^EPIC^ (red), which we expect to adopt a more extended conformation than d6.4, elutes earlier than d6.4 despite having identical molecular weight. It is possible that d6.4^EPIC^ forms a dimer at 2 µM, mediated by hydrophobic residues on the open lid, but the low fluorescence anisotropy of the lid-attached dye (Fig. 4B) suggests that lid-mediated dimerization is unlikely at 75 nM (the conditions of the fluorescence polarization assay).

**Fig. S4.**
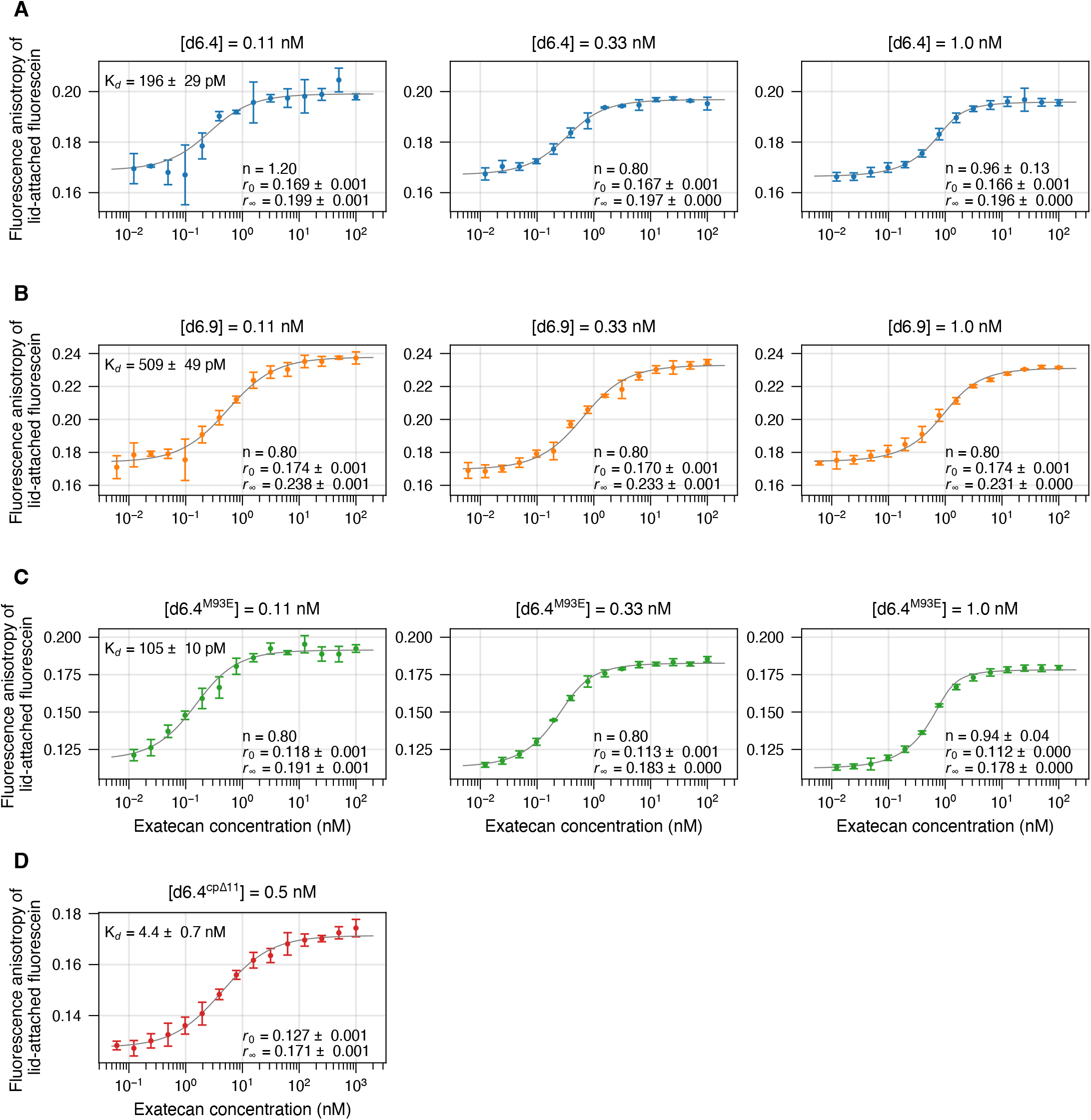
Global fit of binding affinities. Binding affinities of (**A**) d6.4, (**B**) d6.9, (**C**) d6.4^M93E^, and (**D**) d6.4^cpΔ11^ are determined from a global fit across multiple protein concentrations (0.11 nM, 0.33 nM, and 1.0 nM) using lid-attached fluorescein to monitor the fraction of bound protein as a function of titrated exatecan. Error bars represent the standard deviation of 4 technical replicates. The parameters that minimize the weighted least-squares objective are reported with standard errors (see Materials and Methods). K_d_ is binding affinity, r_0_ is anisotropy of unbound protein, r_∞_ is anisotropy of bound protein, and n_1_, n_2_, n_3_ are floating stoichiometry parameters to account for concentration inaccuracies in each condition, respectively (constrained to 0.8 < n < 1.2; standard errors not reported if n falls on the boundary constraint).

**Fig. S5.**
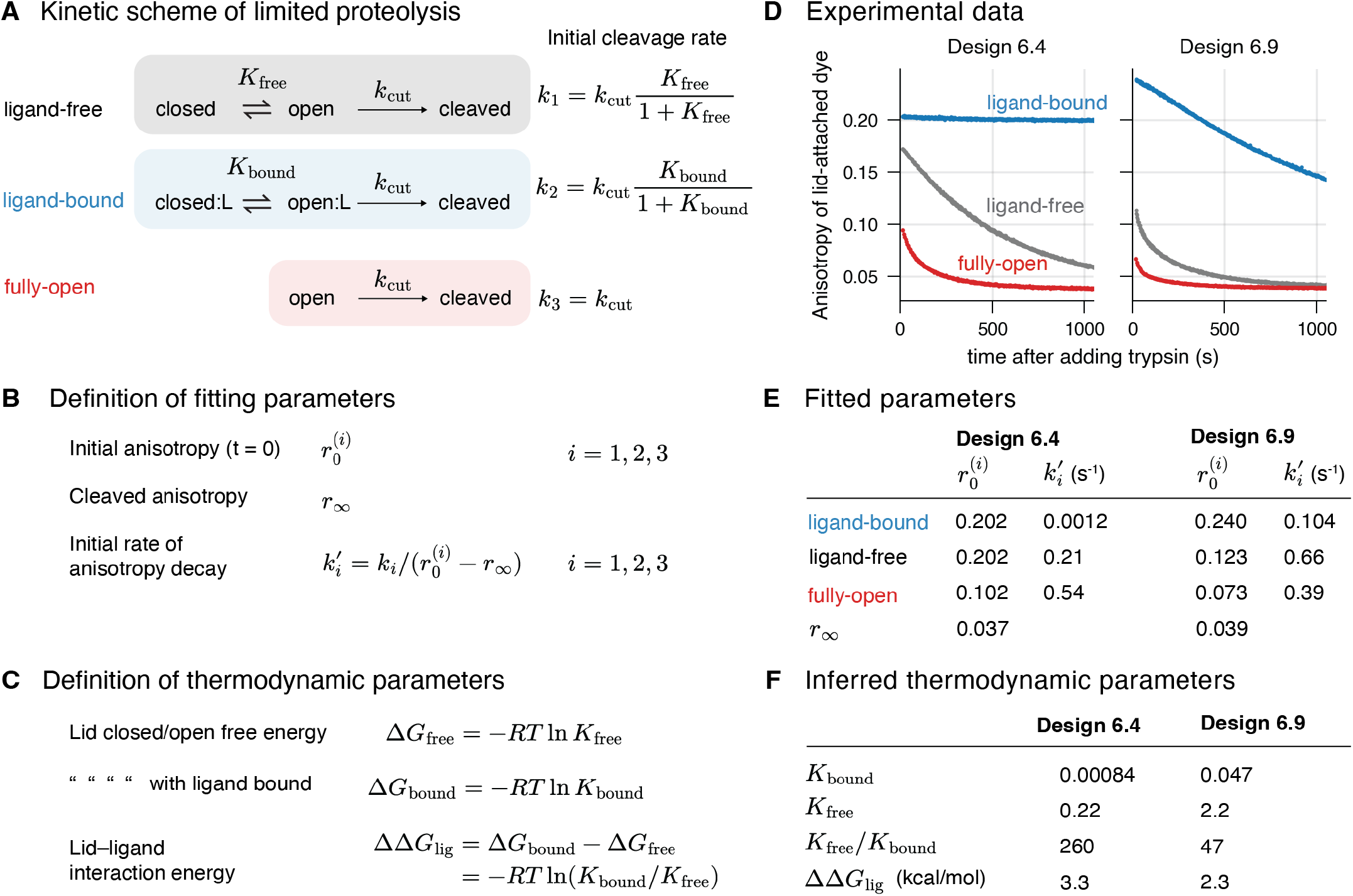
Inferring thermodynamic parameters for d6.4 and d6.9 using limited proteolysis. (**A**) Kinetic scheme for describing proteolytic cleavage of a lid that exchanges between open and closed conformations, assuming that the open state is susceptible to proteolysis with an intrinsic cleavage rate *k*_*cut*_. In the ligand-free and ligand-bound forms, the initial cleavage rates (*k*_1_ and *k*_2_, respectively) depend on the equilibrium fraction in the open state (supplementary text S3). A separate fully-open control protein (see Fig. 4A caption) is used to determine the intrinsic cleavage rate, *k*_3_ = *k*_*cut*_. (**B**) Definition of fitting parameters used to analyze anisotropy decay traces. Each of the three states (open, closed, cleaved) corresponds to a different anisotropy value of a lid-attached fluorescein. The initial anisotropy 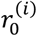 and initial decay rate 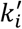 are fitted separately for each of the three conditions (ligand-free, ligand-bound, and fully-open). The final anisotropy *r*_∞_ corresponds to a completely cleaved lid and is common to all three conditions. (**C**) Thermodynamic relationships linking the fitted kinetic parameters to lid conformational equilibria (*K*_free_ and *K*_bound_) and the lid–ligand interaction free energy (ΔΔ*G*_lig_). (**D**) Kinetic traces of proteolysis for d6.4 and d6.9, as monitored through the anisotropy of lid-attached fluorescein. Traces were collected for the ligand-bound state (blue), the ligand-free state (gray), and the fully-open control protein (red). See Materials and Methods for experimental details. (**E**) Fitted kinetic parameters from the experimental data. The parameters 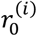 and 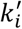 are determined from the slope and intercept, respectively, of a least-squares fit to the initial linear portion of the decay curve, and the parameter *r*_∞_ is taken from the limiting value of the fully-open condition. (**F**) Thermodynamic parameters inferred from the kinetic model, showing that ligand binding shifts the lid-opening equilibrium toward the closed state and stabilizes the closed conformation by 2–3 kcal/mol in d6.4 and d6.9.

**Fig. S6.**
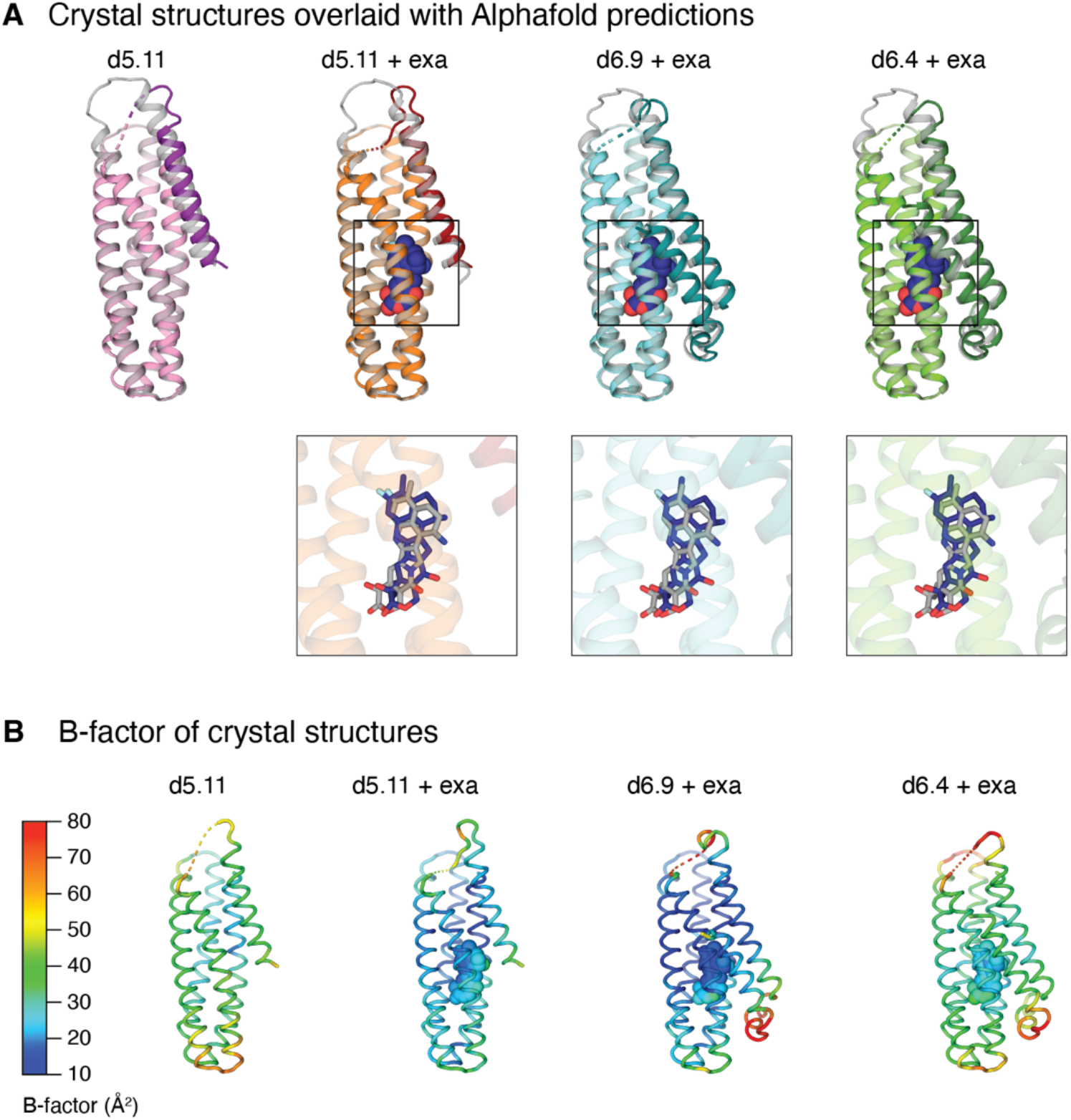
Crystal structures of d5.11, d6.4, and d6.9. (**A**) Crystal structures of the indicated designs (colored) overlaid with the corresponding AlphaFold3 predictions (gray). Insets show the crystallographic pose of the ligand (blue) compared to the AlphaFold3 prediction (gray; ligand heavy-atom RMSD ∼1 Å). The solvent-exposed surface area of the ligand is 10 Å^2^ in the crystal structure of exatecan-bound d5.11 and 0 Å^2^ in d6.9 and d6.4. Design d5.11 crystallized under the same buffer condition both with and without exatecan (table S6). In both cases, the crystal structure shows the same closed, helical conformation of the lid as the design model; however, in the exatecan-free form (leftmost structure in gray), AlphaFold3 incorrectly predicts a register-shifted lid (lid C*α* RMSD = 4 Å, table S3). (**C**) Crystal structures colored by the refined C*α* B-factor. In d6.4 and d6.9, the segment between the two lid helices has the highest B-factors, and the flexible GGS linker is largely unresolved (shown as dashed lines).

**Fig. S7.**
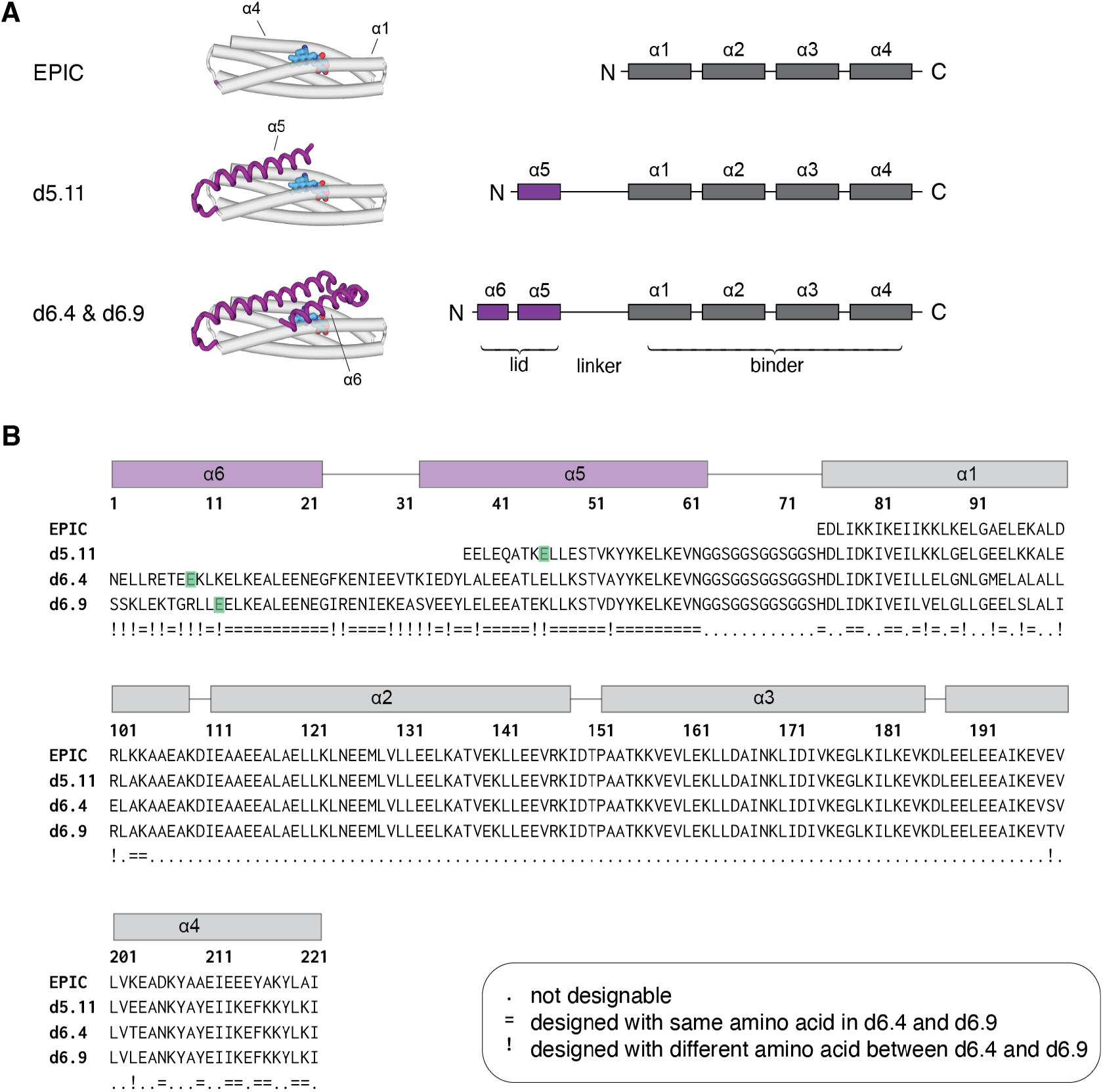
Sequence comparison of EPIC, d5.11, d6.4, and d6.9. (**A**) Topology of the lid domain and binder domain in EPIC, d5.11, d6.4, and d6.9. The purple boxes denote the *α*-helices in the lid domain, and the four gray boxes denote the four *α*-helices of the binder domain. All designs use the sequential helix numbering of *α*1–4 from EPIC. (**B**) Sequence alignment of EPIC, d5.11, d6.4, and d6.9. Out of the 90 designable positions in d6.4 and d6.9 (marked with . symbol), 29 positions were designed with a different amino acid in d6.4 and d6.9 (marked with ! symbol), and 61 positions with the same amino acid in d6.4 and d6.9 (marked with = symbol). The position of the cysteine residue for covalent attachment of fluorescein is highlighted in green.

**Fig. S8.**
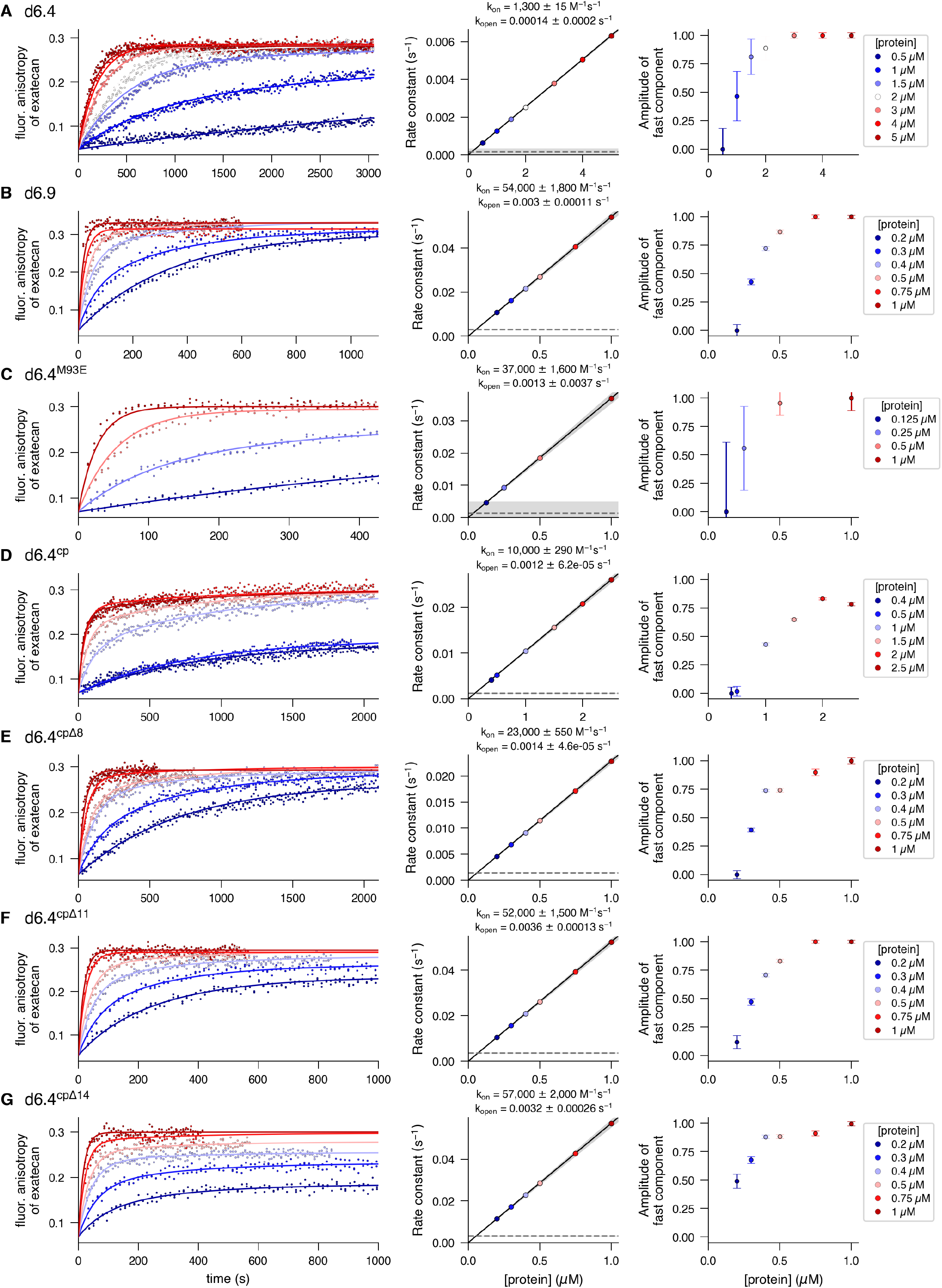
Association kinetics monitored through polarization of intrinsic exatecan fluorescence. Association kinetics of exatecan to (**A**) d6.4, (**B**) d6.9, and (**C**) d6.4^M93E^, (**D**) d6.4^cp^, (**E**) d6.4^cpΔ8^, (**F**) d6.4^cpΔ11^, and (**F**) d6.4^cpΔ14^. Exatecan at 25 nM was mixed with an excess of protein across several protein concentrations (0.5–5 µM for d6.4, 0.5–2 µM for d6.4^cp^, and 0.2–1µM for all others), and the fraction of exatecan bound to protein was monitored over time via fluorescence anisotropy of the ligand’s intrinsic fluorescence. Left panel: the kinetic traces are fitted to a biexponential function where the rate of the faster component is linear in [P] (with proportionality constant *k*_*on*_, the effective bimolecular rate constant) and the slower component is [P]-independent (*k*_*open*_, the lid opening rate). These two parameters are fitted globally across protein concentrations, and the amplitude of the fast component (*f*) is determined separately for each concentration (see Materials and Methods for details). Middle panel: the fitted value of *k*_*on*_ is shown as the slope of the gray line, and *k*_*open*_ as the horizontal gray line, with the shaded gray region representing one standard error (middle panel). Right panel: the fitted value of *f* for each protein concentration is plotted, with error bars representing one standard error (right panel). At low protein concentrations, the slower, [P]-independent component dominates because the accumulation of a lid-open subpopulation becomes the rate-limiting step for ligand binding. At higher concentration, the faster component, which is proportional to [P], dominates and the amplitude of the fast component, *f*, approaches 1. Throughout the text, *k*_*on*_ refers to the faster component that is linear in [P].

**Fig. S9.**
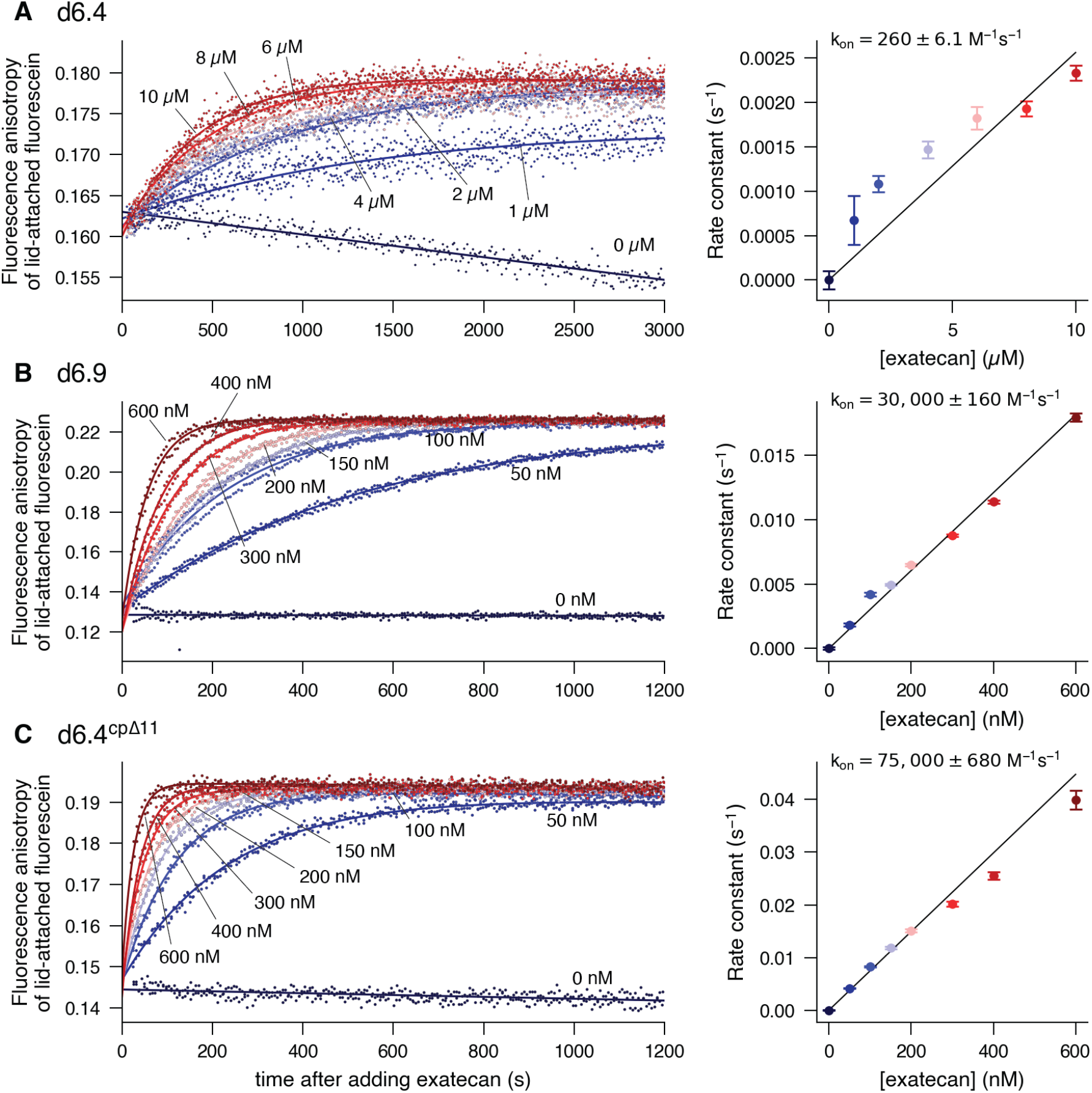
Association kinetics monitored through fluorescence anisotropy of lid-attached dye. Association kinetics of exatecan to (**A**) d6.4, (**B**) d6.9, and (**C**) d6.4^cpΔ11^. A variable amount of exatecan (0–10 µM for d6.4, 0–600 nM for d6.9 and d6.4^cpΔ11^) is added to 10 nM of fluorescein-labeled protein, and the fraction of protein bound to exatecan is monitored as a function of time via the fluorescence anisotropy of the fluorescein attached to the lid. For each concentration of exatecan, a pseudo-first-order rate constant is fitted to an exponential curve with a linear term to account for instrument drift (left panel). The overall second-order rate constant is derived from the slope of a best-fit line across the rates determined at various concentrations (right panel, see Materials and Methods). Error bars represent the standard error of the fitted pseudo-first-order rate constants.

**Fig. S10.**
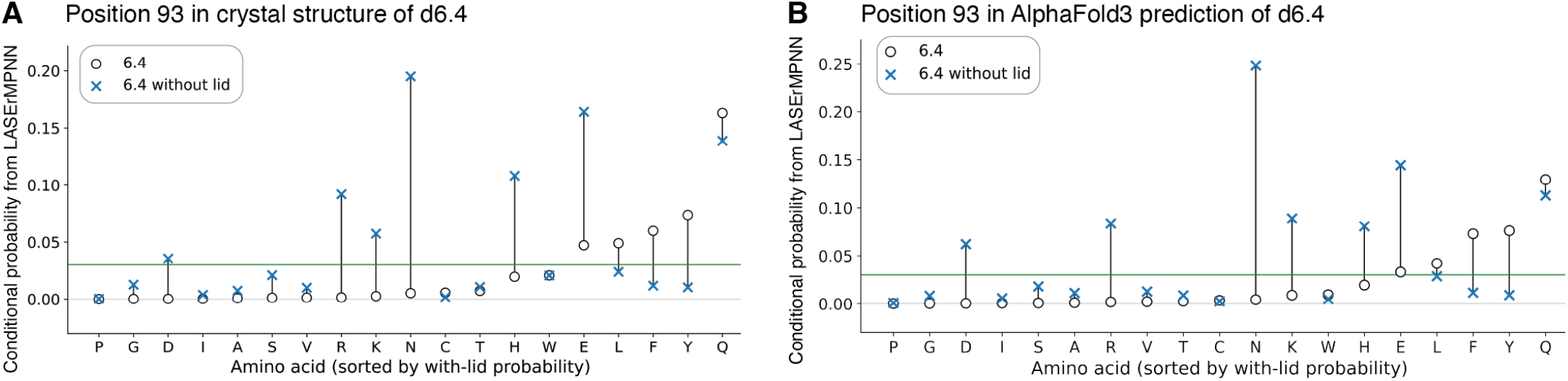
LASErMPNN amino-acid probabilities of position 93 in d6.4. (**A**) To identify suitable mutations for destabilizing the lid–binder interface in d6.4, we calculated the LASErMPNN probabilities of each amino acid at position 93 (a residue on the binder domain in the lid–binder interface), conditioned on the backbone structure, the ligand coordinates, and the residue identities (and side-chain conformations) at every other position. We compared the probabilities for a “closed” structure and an “open” structure, where the closed structure is (**A**) the crystal structure or (**B**) the AF3-predicted structure of d6.4 with exatecan bound, and the open structure is the same structure with the lid removed. The circles show the normalized probabilities in the closed state and crosses in the open state. The wild-type residue M (Met) is omitted from the plot (Prob[position 93 = M] is 53% in closed state, 7% in open state). Amino acids R, K, N, H, and E are predicted with a higher probability in the open than the closed state; of these proposed substitutions, E is the only amino acid that remains compatible with a closed state (> 3% probability, green line).

**Fig. S11.**
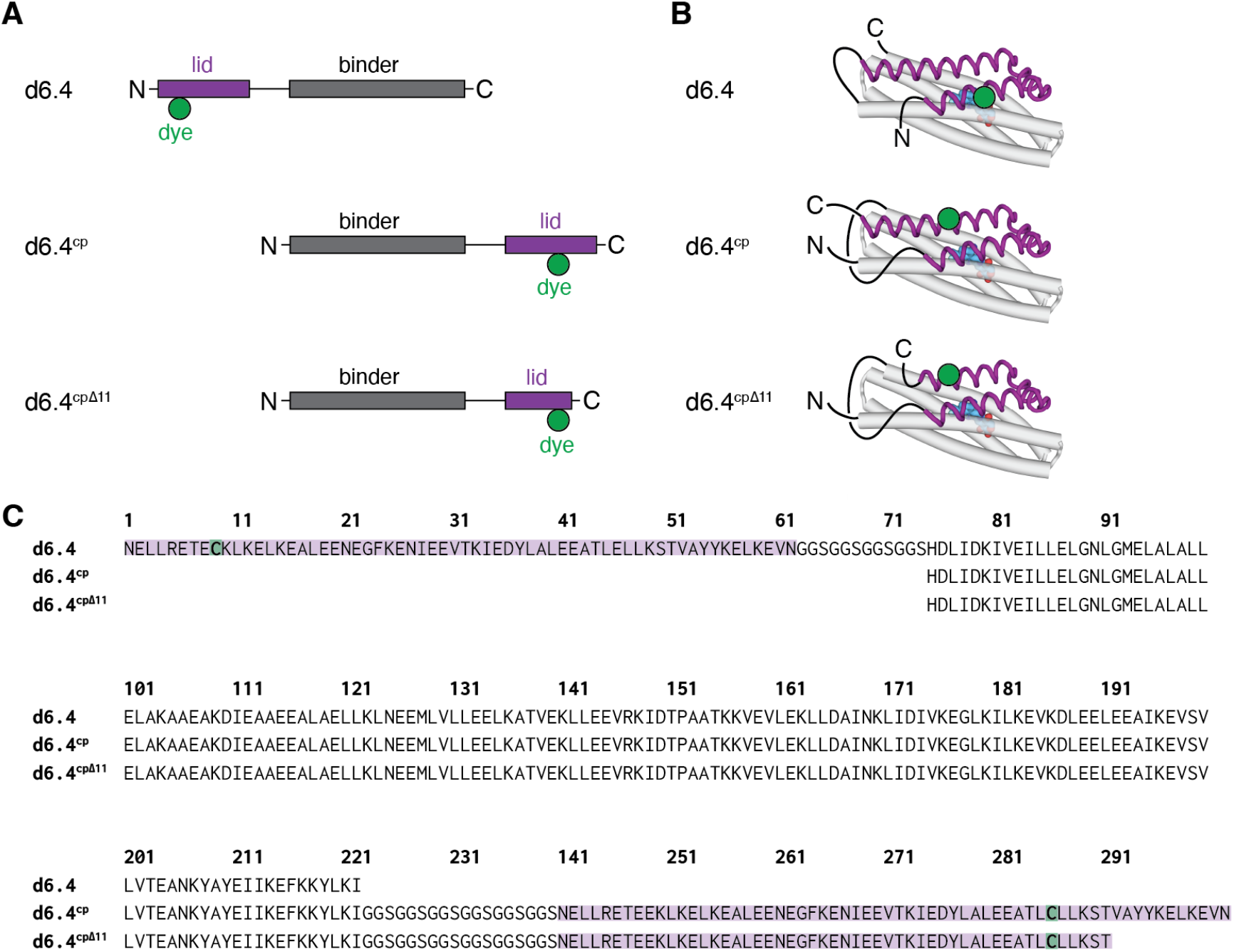
Topology of the circularly permuted protein d6.4^cpΔ11^. (**A**) Topology of d6.4, d6.4^cp^, and d6.4^cpΔ11^. (**B**) Locations of the N- and C-termini in the three-dimensional structure of d6.4, d6.4^cp^, and d6.4^cpΔ11^. The circular permutation in d6.4^cp^ brings the termini closer together to potentially maximize the FRET efficiency between donor and acceptor fused to the termini. Furthermore, in the circularly permutated topology, the C-terminus is positioned further from the ligand (blue) so that truncation does not interfere with lid–ligand interactions. The green circle indicates the position of the solvent-exposed cysteine residue that is used to covalently conjugate a fluorescein to the lid. In the circularly permuted topology, fluorescein is moved to a position closer to the terminus to maximize the difference in anisotropy between the open and closed states (supplementary text S2). (**C**) Sequence alignment of the three sequences, corresponding to the schematics in **A**. The lid portion is highlighted in purple and the fluorescein attachment point is highlighted in green.

**Fig. S12.**
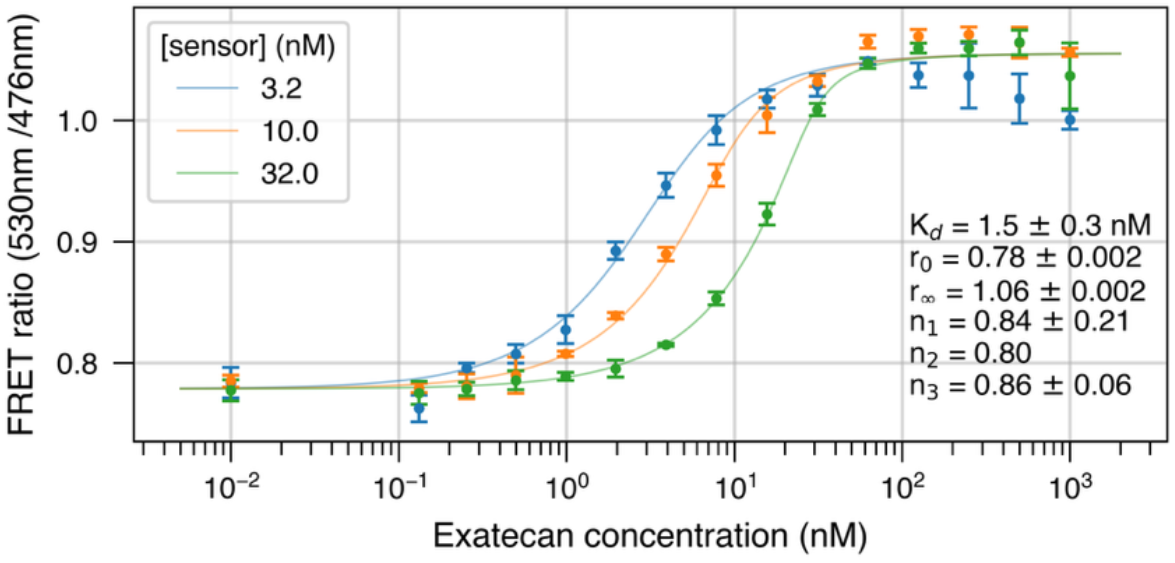
Global fit of binding affinity for FRET sensor mT-d6.4^cpΔ11^-mV. Titration of exatecan into PBS pH 6.5 with the sensor at 3.2, 10.0, and 32 nM. The binding affinity is fit globally across all three sensor concentrations using a quadratic binding model that accounts for depletion of free ligand by sensor binding (see Materials and Methods for definitions). Error bars represent the standard deviation of 3 technical replicates. The parameters that minimize the weighted least-squares objective are reported with standard errors (Materials and Methods). K_d_ is binding affinity, r_0_ is FRET ratio of unbound sensor, r_∞_ is FRET ratio of bound sensor, and n_1_, n_2_, n_3_ are floating stoichiometry parameters to account for concentration inaccuracies under each condition (constrained to 0.8 < n < 1.2; standard error not reported for n_2_ since it falls on the boundary constraint).

**Fig. S13.**
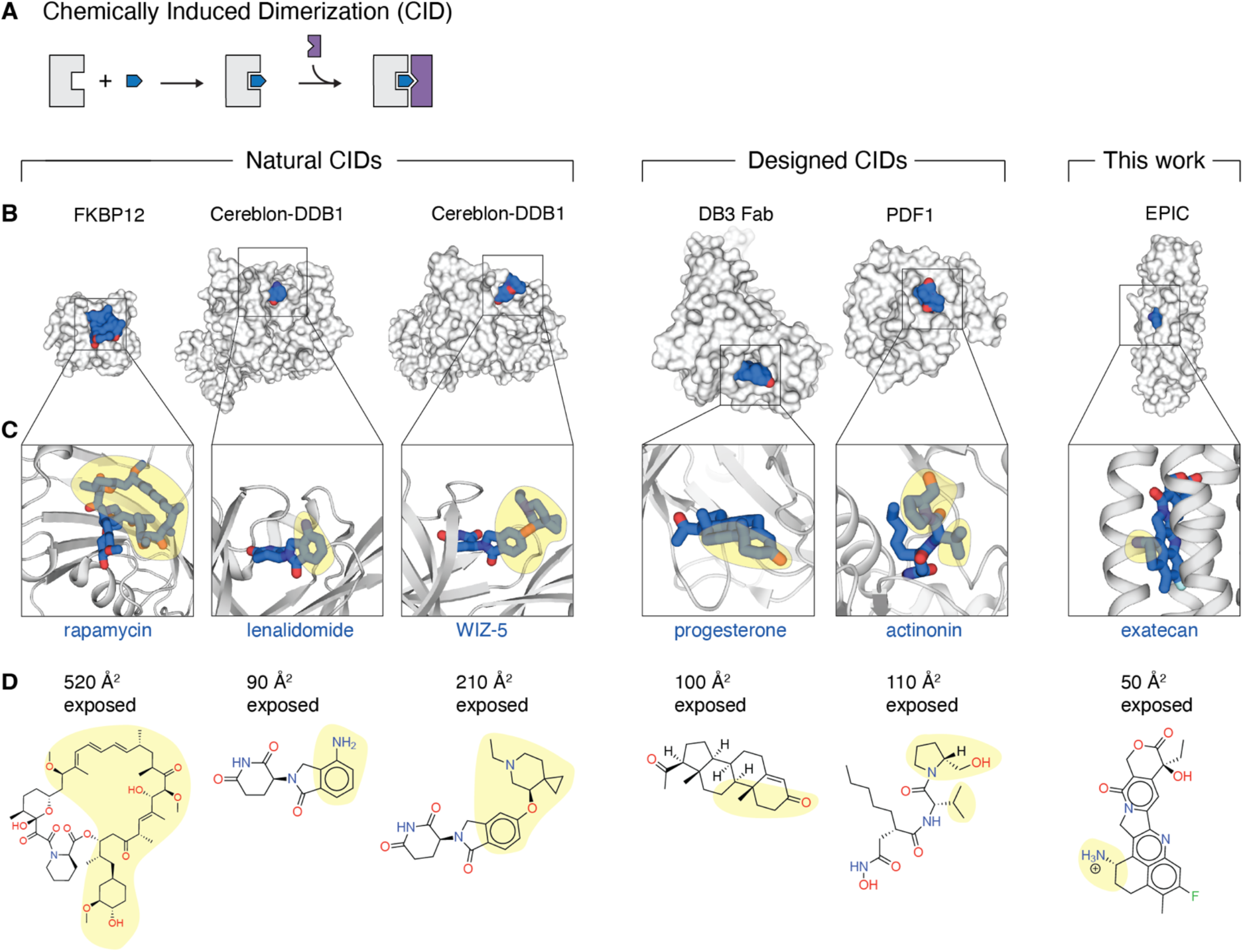
Comparison of exposed ligand surface area for known chemically-induced dimerizers and EPIC. (**A**) For chemically induced dimerizers (CIDs), a small molecule L (blue) first binds one protein partner A (gray), and the A-L complex subsequently binds the second protein B (purple). Protein B selectively binds the A-L complex over A alone because it recognizes the exposed portion of L in the A-L complex. The larger the exposed surface of L, the easier it is for B to distinguish between A and A-L. (**B**) A surface representation of the A-L complex in several examples of known CIDs, with the protein surface in white and the exposed ligand surface in blue. “Natural CIDs” refers to systems where both A and B are naturally occurring proteins; “designed CIDs” refers to ones where A is natural but B is designed. “This work” shows the starting point for the lid design discussed in this work. PDB accession codes: 5GPG, 4CI2, 9DJT, 9FK_d_, 8S1X, and 9NZG, respectively. (**C**) A close-up view of the ligand in the A-L complex, with the solvent-exposed portion of the ligand highlighted in yellow. (**D**) Chemical structures of each of the ligands, again with the exposed portion highlighted in yellow. The solvent-accessible surface area of the ligand (Å^2^) was calculated using freesasa. In many examples of CIDs, the exposed ligand surface is fairly large and predominantly hydrophobic, which simplifies the recognition task—burial of a hydrophobic surface is favorable and feasible as long as the interaction partner has a generally complementary shape. By contrast, our work makes use of an epitope that is both smaller and more polar, which is challenging since it requires precisely positioned interactions to derive sufficient thermodynamic driving force (supplementary text S5).

**Fig. S14.**
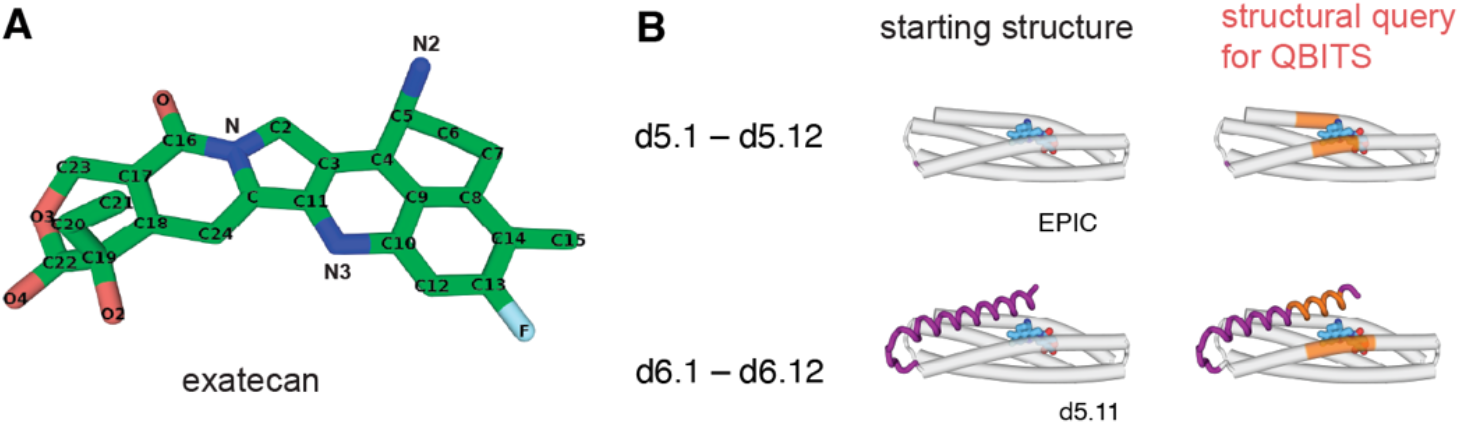
Details of computational pipeline. (**A**) Name of each atom in exatecan. (**B**) Structural query for QBITS used in the design of 5-helix proteins (single-helix lids) and 6-helix proteins (two-helix lids). Residue numbers are 86–93 and 206–213 (for 5-helix proteins) and 40–49 and 85–96 (for 6-helix proteins). Residue numbering is defined in fig. S7. See Materials and Methods for details.

## Supplementary Tables

**Table S1.** Summary of computational workflow for 5-helix designs, d5.1–d5.12..

| Step of Workflow | Number | Comments |
| --- | --- | --- |
| # initial placements of $\alpha 5$ helix | 250 | |
| ... after salt bridge filter | 58 |  |
| ... after crossing angle filter | 45 |  |
|  |  | ×12 helical extensions per placement |
| # templates after helix extension | 540 |  |
|  |  | ×4 weights: [0.0, 0.5, 0.8, 1.0]<br>×2 salt bridge fixed: [True, False]<br>×10 random sequence samples |
| # sequences designed | 43,200 |  |
| ...AF3 drug-bound RMSD < 3 Å and pAE < 5 | 543 |  |
| ...AF3 drug-free RMSD > 3 Å and pAE > 15 | 25 |  |
| ... selected for experimental testing | 12 |  |

**Table S2.**
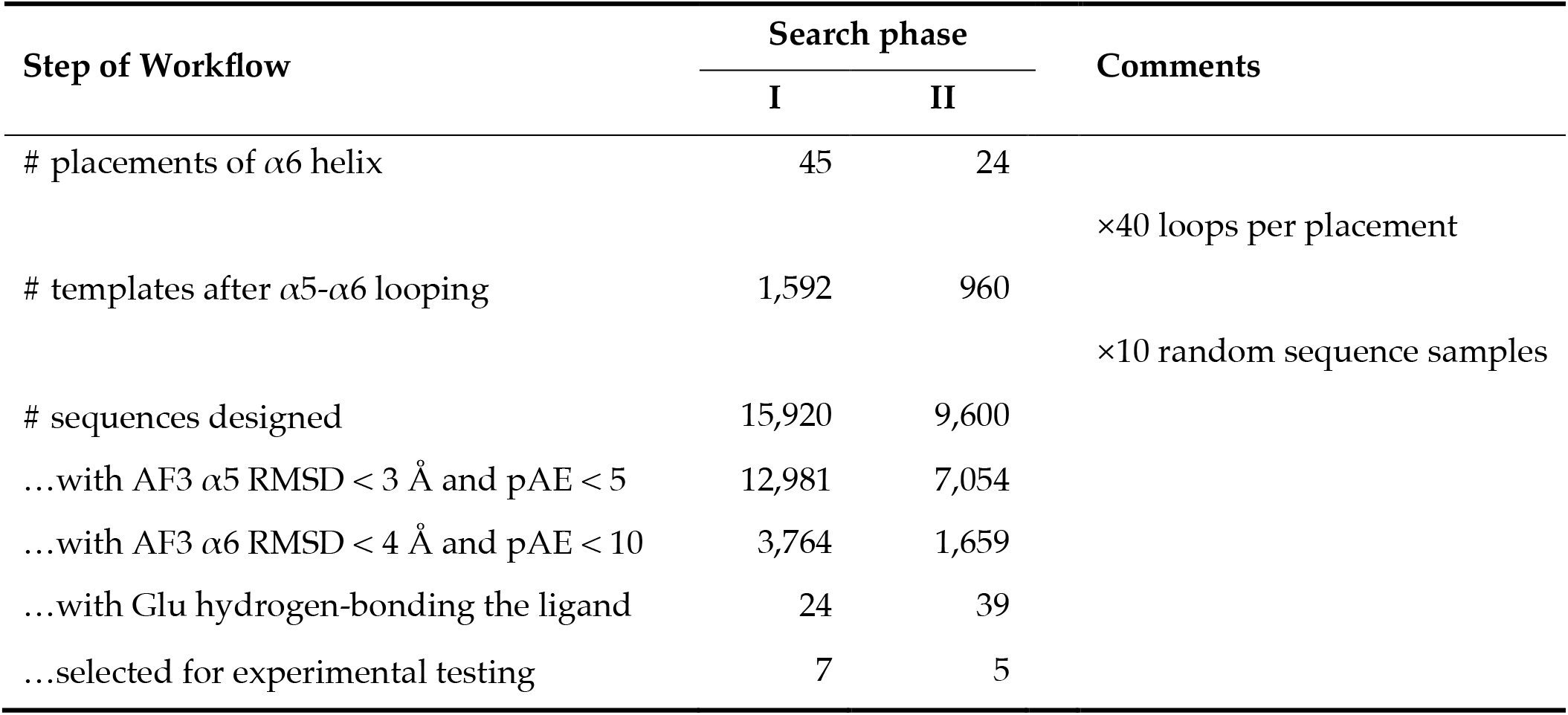
Summary of computational workflow for 6-helix designs, d6.1–d6.12.

**Table S3.** Summary of computational metrics across 24 designs.

| Design | AF3 iPAE of lid |  | AF3 RMSD of lid |  |
| --- | --- | --- | --- | --- |
|  | – exa | + exa | – exa | + exa |
| d5.1 | 30.9 | 4.2 | 96.2 | 2.8 |
| d5.2 | 19.9 | 4.7 | 40.8 | 2.7 |
| d5.3 | 16.7 | 4.7 | 19.1 | 2.5 |
| d5.4 | 24.3 | 4.0 | 14.1 | 2.7 |
| d5.5 | 18.8 | 4.1 | 7.9 | 1.8 |
| d5.6 | 18.4 | 3.0 | 6.5 | 1.8 |
| d5.7 | 23.2 | 4.3 | 6.4 | 2.1 |
| d5.8 | 15.6 | 4.6 | 6.1 | 1.5 |
| d5.9 | 20.3 | 5.0 | 4.8 | 2.3 |
| d5.10 | 22.3 | 4.3 | 4.6 | 1.9 |
| d5.11 | 15.6 | 4.2 | 4.0 | 1.0 |
| d5.12 | 15.4 | 3.3 | 3.4 | 1.7 |
| d6.1 | 5.7 | 5.2 | 1.1 | 0.8 |
| d6.2 | 11.5 | 3.5 | 13.1 | 1.0 |
| d6.3 | 4.6 | 3.1 | 1.1 | 1.1 |
| d6.4 | 14.4 | 4.1 | 18.6 | 1.1 |
| d6.5 | 4.6 | 5.2 | 1.0 | 1.3 |
| d6.6 | 7.9 | 4.6 | 1.3 | 1.4 |
| d6.7 | 8.0 | 4.3 | 1.6 | 1.5 |
| d6.8 | 12.7 | 3.2 | 6.6 | 1.7 |
| d6.9 | 10.9 | 6.4 | 13.0 | 1.0 |
| d6.10 | 9.5 | 6.1 | 1.3 | 1.4 |
| d6.11 | 12.8 | 6.8 | 5.6 | 1.6 |
| d6.12 | 3.6 | 3.2 | 1.6 | 1.5 |

**Table S4.** Summary of experimental results across 24 designs.

| Design | Anisotropy of lid-attached fluorescein |  | Proteolytic resistance<br>(1 / initial rate, sec) |  | K<sub>D</sub> (nM) |
| --- | --- | --- | --- | --- | --- |
|  | – exa | + exa | – exa | + exa |  |
| **d5.1** | 0.102 | 0.102 | 160 | 140 | 280 |
| **d5.2** | 0.190 | 0.180 | 290 | 180 | 2 |
| **d5.3** | 0.239 | 0.229 | 550 | 470 | 250 |
| **d5.4** | 0.154 | 0.141 | 1,600 | 1,300 | 930 |
| **d5.5** | 0.148 | 0.138 | < 100 | < 100 | 800 |
| **d5.6** | 0.172 | 0.167 | 5,700 | 2,700 | 360 |
| **d5.7** | 0.106 | 0.109 | < 100 | < 100 | 140 |
| **d5.8** | 0.174 | 0.193 ^ | 830 | 1,000 | 430 |
| **d5.9** | 0.094 | 0.096 | < 100 | < 100 | 110 |
| **d5.10** | 0.193 | 0.194 | 140 | 180 | 180 |
| **d5.11** | 0.227 | 0.222 | 6,000 | 16,000 * | 6 |
| **d5.12** | 0.207 | 0.199 | 550 | 290 | N.D. |
| **d6.1** | 0.156 | 0.180 ^ | 3,200 | 12,000 * | < 2 |
| **d6.2** | 0.188 | 0.188 | 430 | 21,000 ** | < 2 |
| **d6.3** | 0.218 | 0.219 | > 100,000 | > 100,000 | N.D. |
| **d6.4** | 0.167 | 0.200 ^ | 500 | 78,000 ** | 0.2<sup>†</sup> |
| **d6.5** | 0.132 | 0.130 | 1,600 | 5,000 * | < 2 |
| **d6.6** | 0.116 | 0.129 | < 100 | < 100 | 100 |
| **d6.7** | 0.196 | 0.208 | 260 | 28,000 ** | 3 |
| **d6.8** | 0.123 | 0.127 | < 100 | < 100 | < 2 |
| **d6.9** | 0.131 | 0.224 ^^ | < 100 | 1,300 ** | 0.5<sup>†</sup> |
| **d6.10** | 0.190 | 0.192 | < 100 | < 100 | 20 |
| **d6.11** | 0.172 | 0.179 | 160 | 1,000 * | 4 |
| **d6.12** | 0.194 | 0.197 | 360 | 15,000 ** | < 2 |

^ change in anisotropy > 0.015
^^ change in anisotropy > 0.05
\* ratio in proteolytic rate > 2
\*\* ratio in proteolytic rate > 15
<sup>†</sup> determined through anisotropy of lid-attached fluorescein (Fig. S4)

**Table S5.** Crystallization conditions.

| Design* | MCSG<sup>†</sup> | Mother liquor |
| --- | --- | --- |
| d5.11 | 2/D9 | 1 M lithium chloride, 0.1 M sodium citrate pH 4 (adj. HCl), 20% (w/v) PEG 6000 |
| d5.11 + exa | 2/D9 | 1 M lithium chloride, 0.1 M sodium citrate pH 4 (adj. HCl), 20% (w/v) PEG 6000 |
| d6.9 + exa | 1/D12 | 0.2 M ammonium chloride pH 6.3, 20% (w/v) PEG 3350 |
| d.4 + exa | 1/E7 | 0.2 M ammonium iodide, 20% (w/v) PEG 3350 |

\* exa = exatecan. <sup>†</sup> plate and well number in the MCSG sparse matrix screen.

**Table S6.**
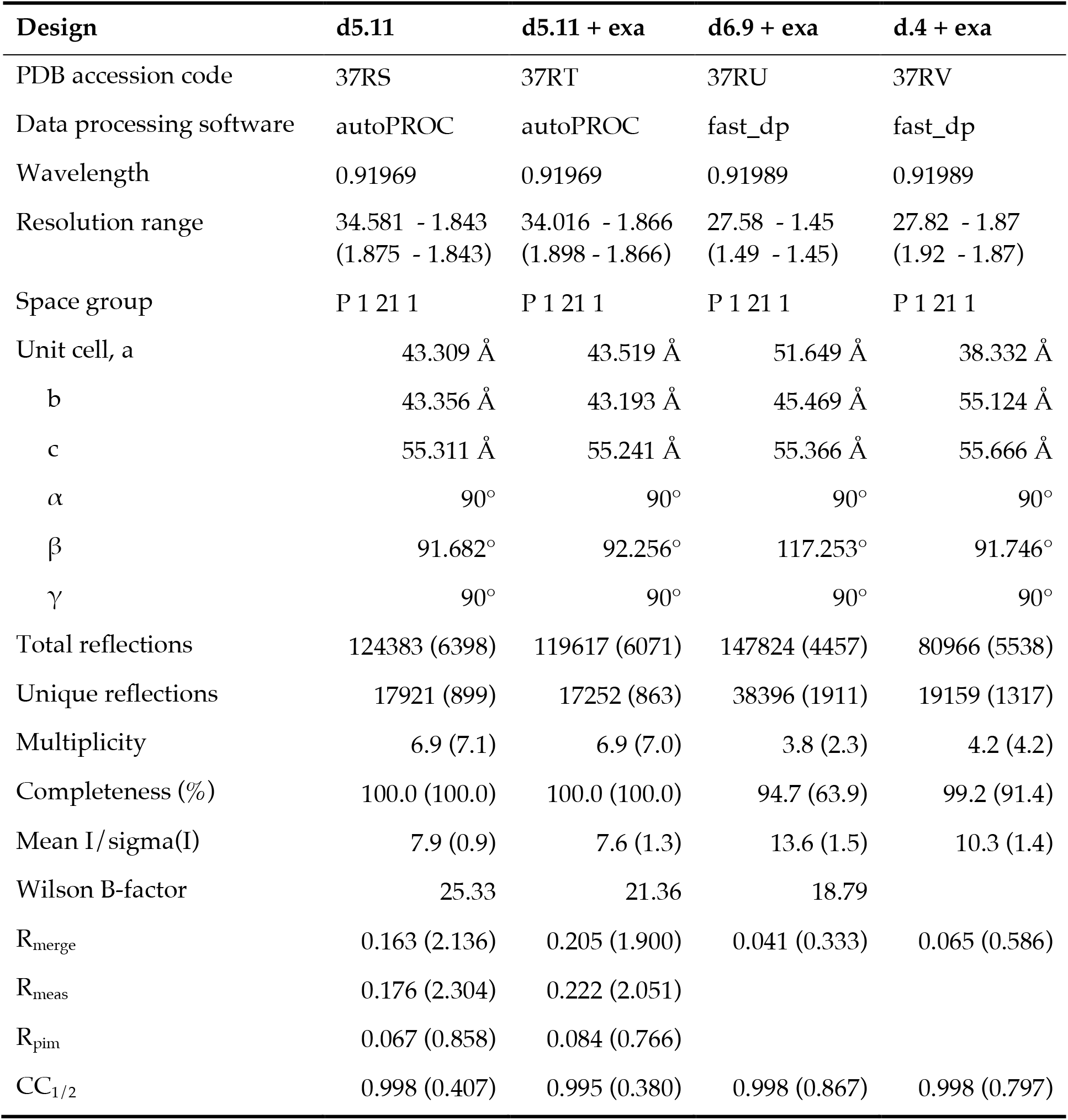
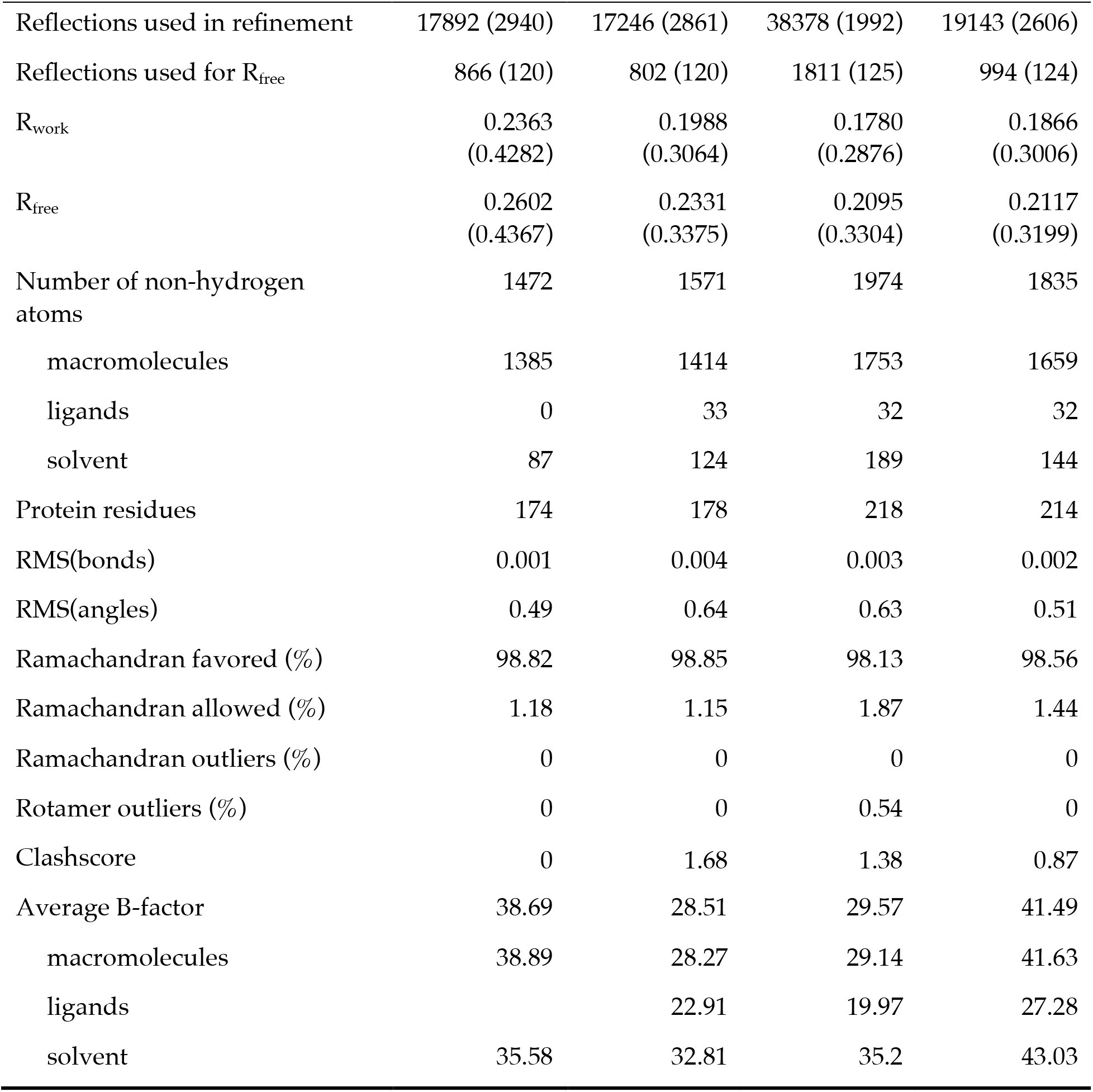
Crystallographic data collection and refinement statistics.

**Table S7.** Summary of experimental results for optimizing design 6.4.

| Design | Anisotropy of lid-attached fluorescein |  | On-rate<sup>†</sup><br>(M<sup>-1</sup> s<sup>-1</sup>) | K<sub>d</sub> (nM) |
| --- | --- | --- | --- | --- |
|  | – exa | + exa |  |  |
| **d6.4** | 0.167 | 0.200 | 1,300 | 0.20 ± 0.03 |
| **d6.4**<sup>M93E</sup> | 0.115 | 0.192 | 37,000 | 0.11 ± 0.01 |
| **d6.4**<sup>EPIC</sup> | 0.104 | 0.104 | > 300,000 | N.D. |
| **d6.4**<sup>cp</sup> | 0.225 | 0.239 | 10,000 | N.D. |
| **d6.4**<sup>cpΔ8</sup> | 0.171 | 0.220 | 23,000 | N.D. |
| **d6.4**<sup>cpΔ11</sup> | 0.148 | 0.210 | 52,000 | 4.4 ± 0.7 |
| **d6.4**<sup>cpΔ14</sup> | 0.128 | 0.134 | 57,000 | N.D. |

<sup>†</sup> determined from the intrinsic fluorescence of exatecan from a global fit across protein concentrations (Fig. S8)

**Table S8.**
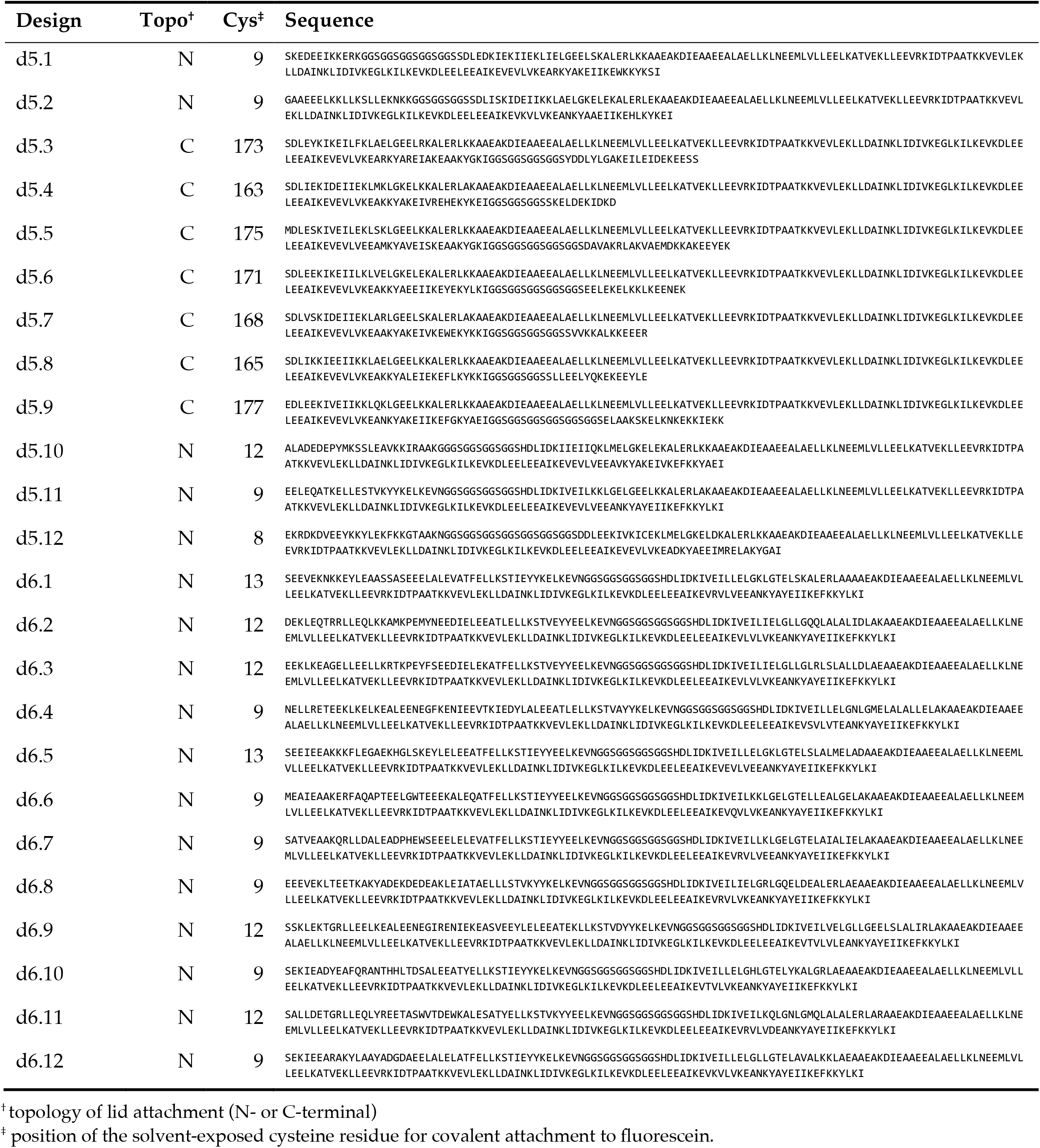
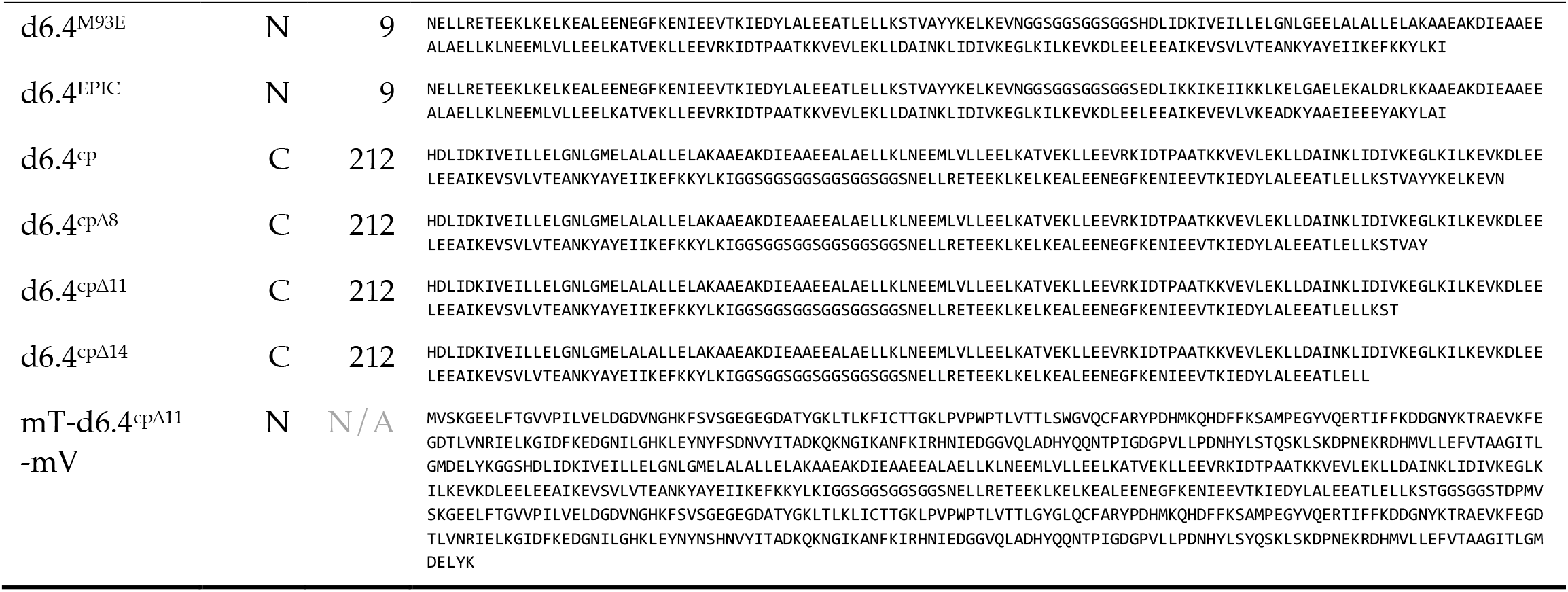
Sequences of proteins in this study.

